# Glycation modulates glutamatergic signalling and exacerbates Parkinson’s disease-like phenotypes

**DOI:** 10.1101/2021.08.24.457507

**Authors:** Ana Chegão, Mariana Guarda, Bruno M. Alexandre, Liana Shvachiy, Mariana Temido-Ferreira, Inês Marques-Morgado, Bárbara Fernandes Gomes, Rune Matthiesen, Luísa V. Lopes, Pedro R. Florindo, Ricardo Anjos-Gomes, Patrícia Gomes-Alves, Joana E. Coelho, Tiago Fleming Outeiro, Hugo Vicente Miranda

**Author notes:** Correspondence to: Hugo Vicente Miranda, DysBrainD: Dysmetabolism in Brain Diseases Lab, CEDOC, NOVA Medical School, Universidade NOVA de Lisboa, Rua Câmara Pestana, № 6, Edifício CEDOC II, Office 2.27, 1150-082 Lisboa, Portugal, Correspondence may also be addressed to: Tiago Fleming Outeiro, Department of Experimental Neurodegeneration, University Medical Center Gottingen, Waldweg 33, 37073 Gottingen, Germany.

## Abstract

Alpha-synuclein (aSyn) is assumed to be a central player in the pathogenesis of synucleinopathies due to its accumulation in typical protein aggregates in the brain. However, it is still unclear how it contributes to neurodegeneration. Type-2 diabetes mellitus is a risk factor for Parkinson’s disease and, one common molecular alteration among these disorders is an age-associated increase in protein glycation. Thus, we hypothesized that glycation-induced dysfunction of neuronal pathways might be an underlying molecular cause of synucleinopathies. Here, we evaluated if increased brain glycation modulated motor and/or non-motor phenotypes in a mouse model of synucleinopathies. In addition, we dissected the specific impact of methylglyoxal (MGO, a glycating agent) in mice overexpressing aSyn in the brain, and unveiled the major molecular pathways altered. Age-matched (16 weeks old) male aSyn transgenic (Thy1-aSyn) or WT mice received a single dose of MGO or vehicle via intracerebroventricular (ICV) injection. Behavioural phenotypes were analysed 4 weeks post-treatment, and, at the end of the tests, biochemical and histological studies were conducted on brain tissue. We found that glycation potentiates motor dysfunction, assessed by vertical pole, rotarod and hindlimb clasping tests in Thy1-aSyn mice. In addition, it induces cognitive impairment (Y maze test), olfactory disturbances (block test), and colonic dysfunction. These behavioural changes were accompanied by the accumulation of aSyn in the midbrain, striatum, and prefrontal cortex, and by an overall increase in glycation in the midbrain and cerebellum. Furthermore, MGO induced neuronal and dopaminergic cell loss in the midbrain of Thy1-aSyn mice. Quantitative proteomic analysis revealed that, in Thy1-aSyn mice, MGO mainly impacts on glutamatergic proteins in the midbrain, but not in the prefrontal cortex, where it mainly affects the electron transport chain. Among the altered proteins in the midbrain, we found an upregulation of N-Methyl-_D_-Aspartate (NMDA) and α-amino-3-hydroxy-5-methyl-4-isoxazolepropionic acid (AMPA) glutamate receptors levels, glutaminase, vesicle glutamate transporter (VGLUT), and the excitatory amino acid transporter (EAAT1), suggesting potentiation of glutamatergic signalling. Overall, we demonstrated that MGO-induced glycation accelerates Parkinsonian-like sensorimotor and cognitive alterations. The increase in glutamatergic-related proteins in the midbrain may represent a compensatory mechanism to the MGO-induced dopaminergic neurodegeneration. Our study sheds light into the enhanced vulnerability of the midbrain in Parkinson’s disease-related synaptic dysfunction that, ultimately leads to cell loss, and provides molecular insight into the observation that glycation suppressors and anti-glutamatergic drugs hold promise as disease-modifying therapies for synucleinopathies.

## Introduction

Aging is an inevitable process that increases the risk for a number of conditions, including neurodegenerative disorders, such as Parkinson’s disease. Neurodegenerative disorders are typically associated with the misfolding and aggregation of specific proteins in the brain and other tissues. However, the molecular mechanisms that trigger these phenomena are still elusive.

Parkinson’s disease is one of several disorders known as synucleinopathies due to the misfolding and aggregation of alpha-synuclein (aSyn)^1,2^, a protein abundant in the brain that is also present in other tissues. Synucleinopathies include dementia with Lewy bodies, multiple system atrophy, and pure autonomic failure^3–5^. Parkinsonism is a clinical syndrome characterized by resting tremor, bradykinesia, muscular rigidity, postural instability, and gait impairment^6–8^. Nevertheless, the clinical spectrum of Parkinson’s disease includes several non-motor features such as cognitive impairment, hyposmia, obstipation, anxiety and depression, sleep disturbances, pain, and fatigue ^7,9–11^.

aSyn is a natively unfolded protein that, under certain conditions, is prone to aggregation^2,12–15^. Several factors contribute to the oligomerization and fibrillization of aSyn, including high protein concentration, molecular crowding, mutations, posttranslational modifications (PTMs), and interactions with specific metals and small molecules^16,17^.

Physiologically, Parkinson’s disease is characterized by pronounced synaptic alterations and dysregulation of multiple neurotransmission pathways. Pathologically, Parkinson’s disease is characterized by the loss of dopaminergic neurons in the substantia nigra pars compacta (SNpc) and by the presence of proteinaceous aggregates primarily composed of aSyn, known as Lewy bodies and Lewy neurites, in surviving neurons^1,18–21^. The pathological oligomerization and aggregation of aSyn is suggested to, somehow, trigger the degeneration of dopaminergic neurons in the SNpc^20–22^ and, the consequent depletion of dopamine in the striatum induces alterations in dopamine signalling^23–26^ and major functional alterations in glutamatergic synapses^27–32^. Glutamate is the predominant excitatory neurotransmitter in the basal ganglia (BG)^33^. Although the striatum contains the highest density of glutamate receptors in the BG, glutamate or glutamate-dopamine neurons are intermixed with midbrain dopaminergic neurons. Interestingly, the SNpc is rich in NMDA and AMPA glutamate receptors, as well as in metabotropic glutamate receptors^34–36^. It is believed that the increase in firing of subthalamic nucleus neurons in Parkinson’s disease acts as a compensatory mechanism to increase the release of dopamine from the surviving dopaminergic neurons in the SNpc in order to maintain dopamine homeostasis^37^. Furthermore, dopaminergic denervation also induces dysfunctional cortico-striatal glutamate release^38^, thereby eliciting an excitotoxic cascade that promotes further dopaminergic neuronal loss and neurodegeneration^39–41^.

Since genetic alterations account for a smaller fraction of Parkinson’s disease cases, it is imperative to identify and better understand the role of risk factors in the pathogenesis of this disorder. Type-2 diabetes mellitus, a widespread chronic metabolic disease, has been established as an important risk factor for Parkinson’s disease and other neurodegenerative diseases^42,43^. Epidemiological studies revealed that 80% of Parkinson’s disease patients have impaired glucose metabolism. Moreover, diabetes can increase the risk of developing Parkinson’s disease in young diabetic individuals by up to 380%, and significantly accelerates the progression of both motor and cognitive deficits in Parkinson’s disease patients^44–46^. However, the molecular mechanisms underlying this correlation are still unclear. One of the major outcomes of type-2 diabetes mellitus is the deleterious accumulation of reducing sugars that are unavoidably formed as by-products of essential metabolic processes such as glycolysis^47–50^. Glycation is a non-enzymatic reaction between reducing sugars, such as methylglyoxal (MGO), and biomolecules such as proteins, leading to the formation of mostly-irreversible advanced glycation end products (AGEs)^49–51^. Importantly, the levels of circulating MGO in type-2 diabetes mellitus patients are 2-4-fold higher than in healthy individuals^52–55^.

Previously, we found that aSyn is glycated in the brains of Parkinson’s disease patients and that this modification exacerbates aSyn pathogenicity by promoting its accumulation, oligomerization, aggregation, and toxicity, in vitro and in vivo^49,56,57^.

In this study, we aimed to determine whether generalized glycation in the mouse brain could trigger Parkinson’s disease-like features, and to identify the molecular pathways implicated in this process. For this, we delivered MGO via ICV injection in transgenic Thy1-aSyn mice and in corresponding littermates, and evaluated behavioural and biological alterations. We found that MGO exacerbates Parkinson’s disease-like motor and non-motor features, alongside with proteomic alterations of glutamatergic components, specifically in the midbrain. In total, our study provides novel mechanistic insight into the connection between metabolic alterations, as those present in type-2 diabetes mellitus, and Parkinson’s disease, opening novel avenues for the design of therapeutic interventions.

## Materials And Methods

### Methylglyoxal production and standardization

MGO was synthesized as previously described^56,58^ and purified by fractional distillation. MGO quantification was performed by its derivatization with aminoguanidine^59,60^ (more details in supplementary material).

### Animals

We started by evaluating if glycation contributes to the onset or if it exacerbates Parkinson’s disease-like features. For that purpose, we used Thy1-aSyn mice as a model of synucleinopathies. This model recapitulates several features of Parkinson’s disease, including aSyn pathology, alterations in nigrostriatal dopaminergic pathway, loss of striatal dopamine and TH, and a progressive neurodegenerative process, with gradual motor and nonmotor deficits^61,62^.

### Mice demographics and groups characterization

The cohort consisted of 27 male Thy1-aSyn and WT littermate mice, distributed among the four experimental groups: vehicle- (1); or MGO-injected (2) WT littermate mice; and vehicle- (3); or MGO-injected (4) Thy1-aSyn mice (Supplementary Fig. 1A and B). Vehicle-injected or MGO-injected WT littermate mice group included 5 animals each, with an average age of 16.00 ± 1.00 weeks and weight of 29.62 ± 4.12 g or 29.01 ± 3.62 g, respectively (Supplementary Fig. 1A and B). Vehicle-injected or MGO-injected Thy1-aSyn mice group was composed of 9 or 8 animals, respectively, with an average age of 16.11 ± 0.78 or 16.25 ± 0.89 weeks and weight of 28.66 ± 1.43 or 28.19 ± 2.47g (Supplementary Fig. 1A and B). Since Thy1-aSyn female mice exhibit much fewer behavioural changes, we decided to only include male mice in our study^61,63^.

### Intracerebroventricular injection of MGO and general procedures

Age-matched 16 weeks old, male transgenic Thy1-aSyn and WT littermates mice received MGO, or vehicle (PBS, pH 7.4) ICV injection under deep anaesthesia (80mg/kg ketamine hydrochloride, 5 mg/kg xylazine hydrochloride). Animals were kept anesthetized using isoflurane (2% – 4%) and kept at a constant body temperature using a conventional heat pad. Briefly, 5 μL of MGO (31.6 mM) or PBS were injected into the right lateral ventricle, with the following stereotaxic coordinates, relative to Bregma: Anterior-Posterior (AP) – 0.5 mm, Medial-Lateral (ML) – −1.0 mm, and Dorsal-Ventral (DV) – −2.0 mm. Behavioural testing was performed starting four weeks after surgery. Upon conclusion of the behavioural phenotyping, mice were sacrificed (Supplementary Fig. 1C).

### Behavioural tests

To evaluate the effects of glycation on behaviour, mice underwent a battery of behaviour tests to characterize motor, cognitive, anxiety-related, and olfactory function. Open field test, pole test, rotarod, wire hang test, and adhesive removal test were performed to evaluate motor function. Y maze test was used to assess spatial short-term memory (hippocampal dependent), as a read-out of cognitive function. Anxiety-related behaviour was evaluated with elevated plus maze test. The block test was performed to assess olfactory function. SHIRPA protocol was also used to evaluate general health behaviour. For behavioural tests details, see Supplementary materials.

### Immunohistochemistry and counting of neuronal populations of mice brain

After mice sacrifice, the right hemisphere of the brain was transferred into paraformaldehyde (PFA) solution for fixation and immunohistological analysis was performed as previously described^56^. Next, they were cryoprotected in TBS (pH 7.6) containing 30% sucrose (w/v) overnight at 4°C. Sagittal free-floating sections (30mm) were cut around the region of substantia nigra using a cryostat (Leica CM 3050S, Germany) and, subsequently stained as reported previously^64^. In short, free-floating sections were blocked (5% Goat Serum (BioWest, France) and 1% Bovine serum (VWR, USA) and then incubated with different primary antibodies, specifically anti-tyrosine hydroxylase (TH) rabbit (Millipore, 1:1000), anti-aSyn (1:1000, BD Transduction laboratories), anti-NeuN mouse (Millipore, 1:400) overnight at 4°C. Sections were then washed with TBS and incubated with Alexa Flour 488/555 secondary antibodies (1:1000, Invitrogen). Sections were mounted in SuperFrost^®^ Microscope Slides using Mowiol mounting media (Calbiochem). Omission of the primary antibody resulted in no staining. Image acquisition (10x for a whole brain; 63x and 100x in the substantia nigra region) was performed under a confocal point-scanning microscope using both z-stack and scan tile (Zeiss LSM 800 with Airyscan).

Numbers of TH-, DAPI- or NeuN-positive neurons were counted manually, blinded for experimental grouping, using Cell Counter plugin to mark cells from Fiji open-source software^65^. For each animal within the groups, at least 4 sagittal slices were quantified, and an average used for statistical analysis.

### Tissue lysate preparation

After sacrifice, mice brains were collected and separated into left and right hemispheres. The left hemisphere was dissected into midbrain, striatum, cerebellum, hippocampus, and prefrontal cortex, and rapidly frozen in liquid nitrogen for biochemical analysis. Tissue lysates were prepared as previously described^56^.

### SWATH-MS analysis

A high-throughput proteomics analysis using NanoLC coupled to the TripleTOF 6600 (at UniMS, Mass Spectrometry Unit at iBET/ITQB) was performed to screen for differences in protein expression between experimental groups of midbrain or prefrontal cortex protein extracts. See supplementary materials for sample preparation, information-dependent acquisition runs to generate the spectral library, protein quantification by SWATH-MS and quantitative analysis.

#### Proteome functional analysis

Venn diagrams of the statistically differently regulated proteins in midbrain or prefrontal cortex between the experimental groups were used to identify the uniquely affected hits affected in Thy1-aSyn mice injected with MGO using the tool InteractiVenn^66^. These were identified via the comparison between Thy1-aSyn mice injected with MGO and vehicle, excluding the common hits between Thy1-aSyn mice injected with vehicle and the WT mice injected with vehicle (excluding hits generally affected by aSyn expression); and between WT mice injected with MGO vs vehicle (excluding hits generally affected by MGO glycation). Representation of the number of statistically significant up and downregulated hits was performed in Graphpad Prism V9 volcano plots. Protein-protein functional associations were retrieved from STRING (http://www.string-db.org/, version 11.0)^67^. We used the online Enrichr tool (http://www.amp.pharm.mssm.edu/Enrichr/) for the functional enrichment analysis, including Kyoto Encyclopaedia of Genes and Genomes (KEGG) and Gene Ontology (GO) resources^68^. Pathways were obtained from “KEGG 2019 Mouse” and GO terms from “GO Biological Process 2018”, “GO Molecular Function 2018”, and “GO Cellular Component 2018”. Top 7 pathways or GO terms with the highest -log10 Fisher exact test p-value were selected (p values bellow 0.05). Heatmaps representing the differently regulated proteins per top 3 KEGG pathways were done in GraphPad Prism version 9. Protein-protein functional associations representations were colour-coded according to the top KEGG and GO analysis in STRING (Search Tool for the Retrieval of Interacting Genes/Proteins)^67,69–78^. The minimum required interaction score was defined for high confidence (0.7) and disconnected nodes in the network were hidden.

### Immunoblot analysis

These procedures were performed as previously. For more details, see extended material and methods. Proteins were probed using N^ε^-carboxyethyl lysine (CEL) (Mouse Anti-N^ε^-carboxyethyl lysine, Cosmo-Bio, USA), aSyn (Purified Mouse Anti-α-Synuclein antibody, BD Biosciences; San Jose, CA, USA), and β-actin (Mouse Monoclonal anti-β-actin antibody Ambion, Thermo Fisher Scientific; Waltham, MA, USA).

### Statistical analysis

Each experimental group was composed at least of five mice, unless stated otherwise, and all values are expressed as normalized means plus standard deviation. Statistical analysis was performed using GraphPad Prism version 9. One-way ANOVA were used to compare differences among conditions and groups, followed by Dunnett’s multiple comparison test. Values of p < 0.05 were considered significant.

## Results

### MGO potentiates motor deficits and accelerates colonic dysfunction in Thy1-aSyn mice

First, in order to establish a baseline, we applied a battery of motor tests in order to characterize the motor behaviour of the animals used in the study (20 weeks-old animals). The open field test was performed to evaluate general motor activity, gross locomotor activity, and exploration habits^79^. The vertical pole test and rotarod were aimed at assessing locomotor activity, motor coordination and balance^79–82^. To evaluate balance and grip strength, the wire hang test was conducted^79,83^. The adhesive removal test was performed to evaluate sensory and motor deficits related to the paw and the mouth^84^. Finally, we also assessed the motor function using SHIRPA protocol tests, mainly the hindlimb clasping test, and colonic function (assessed during open field test)^80,85^.

Vehicle-injected transgenic Thy1-aSyn mice required more time to turn down on the vertical pole (2.1-fold increase) than WT littermates but displayed no significant differences in the time of climbing down or in the total time to perform the task (Fig. 1A-C). Thy1-aSyn mice also showed a decreased latency to fall in the rotarod at both stationary (3.1-fold decrease) or accelerated rotation (1.9-fold decrease) (Fig. 1D and E) when compared to WT littermates. Impairments were also apparent in grip strength evaluated in the wire hang test, with a decreased latency to fall (3.1-fold decrease) (Supplementary Fig. 1D), and in the hindlimb clasping test, showing increased score when compared to WT littermate mice (Fig. 1F). We observed no sensorimotor differences, in the adhesive removal test (Supplementary Fig. 1E), no alterations of locomotor behaviour, assessed by open field (Supplementary Fig. 1F-J), and no differences in colonic function, given by the number of faecal pellets dropped in an open field in 10 minutes (Fig. 1G).

**Figure 1.**
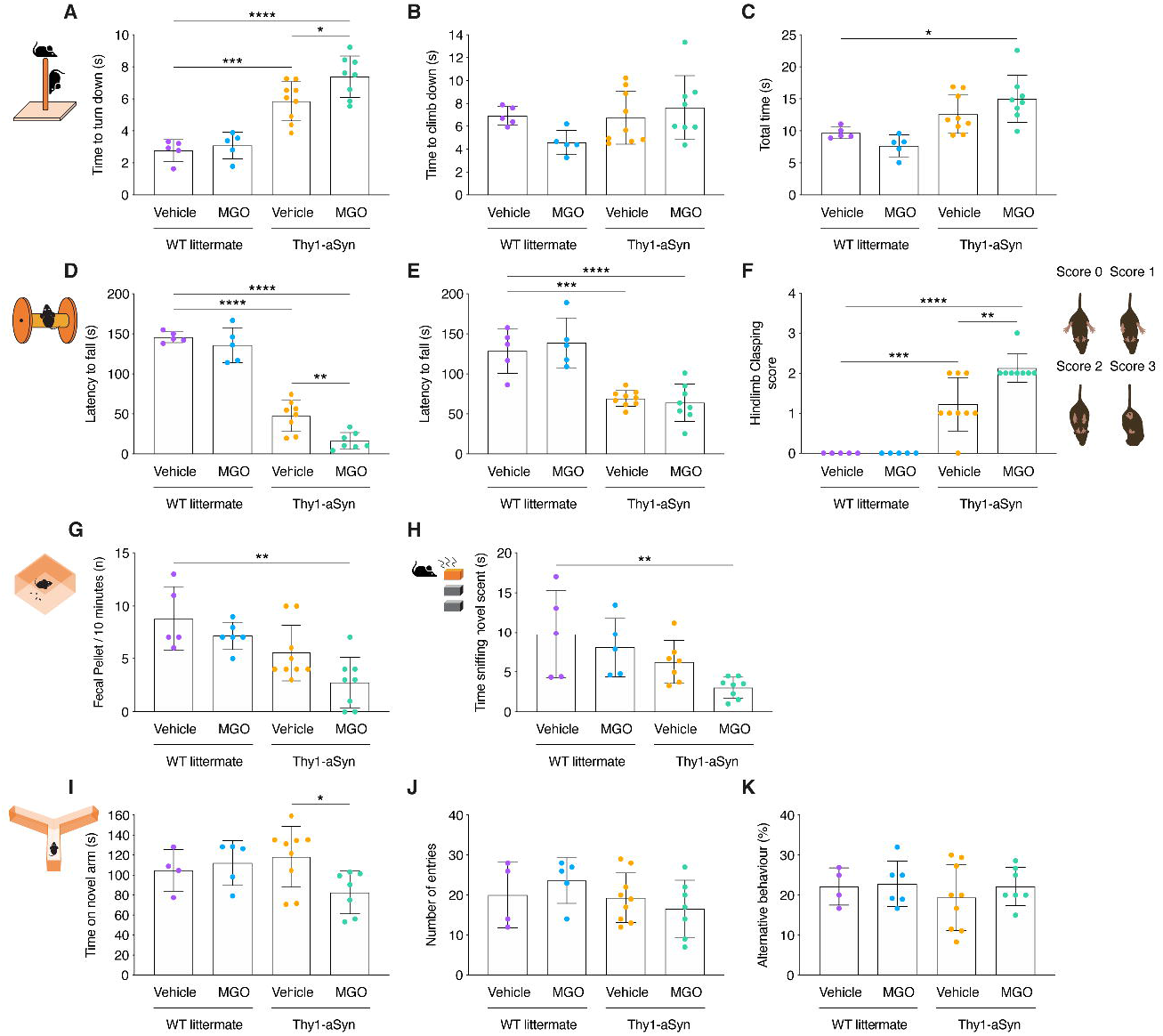
Glycation induces motor dysfunction, cognitive impairment, olfactory, and colonic disturbances in Thy1-aSyn mice. Wild-type littermate (WT) and Thy1-aSyn transgenic (Tg) mice received an intracerebroventricular (ICV) injection of MGO or vehicle (PBS) at 16 weeks of age. Behavioural testing started 4 weeks post-surgery. Plot representations of: pole test - **(A)** time to turn down **(B)** time to climb down, **(C)** total time; rotarod - **(D)** stationary protocol, **(E)** acceleration protocol; **(F)** hindlimb clasping score; **(G)** faecal pellet production; **(H)** block test - time sniffing novel scent; Y maze test - **(I)** time on novel arm, **(J)** number of entries, **(K)** alternative behaviour. At least n=5 in all groups, Ordinary one-way ANOVA, *p < 0.05, **p < 0.01, ***p < 0.001, ****p < 0.0001.

Next, we assessed the effect of MGO injection and found that Thy1-aSyn mice required more time to turn down on the vertical pole compared to vehicle-injected Thy1-aSyn mice (1.3-fold increase), vehicle-injected WT mice (2.7-fold increase) and MGO-injected WT mice (2.4-fold increase) (Fig. 1A). No changes in the time to climb down were observed (Fig. 1B). However, MGO-injected Thy1-aSyn mice took more time to complete this task when compared to vehicle-injected WT mice (1.6-fold increase) or to MGO-injected WT mice (2-fold increase), but not to vehicle-injected Thy1-aSyn mice (Fig. 1C). Rotarod performance was also worse in MGO-injected Thy1-aSyn mice, evaluated at a steady rotation of the rod (2.9-fold decrease of latency to fall), when compared to vehicle-injected Thy1-aSyn mice (Fig. 1D). At accelerated rotation, no differences were observed (Fig. 1E). Finally, MGO-injected Thy1-mice showed worst performance in the hindlimb clasping test (Fig. 1F), and worst colonic function, with a smaller number of faecal pellets, when comparing with vehicle-injected WT mice (Fig. 1G). One the other hand, MGO treatment did not significantly change the grip strength (Supplementary Fig. 1D) sensorimotor function (adhesive test Supplementary Fig. 1E), or locomotor activity, assessed by the open field test (Supplementary Fig. 1F-J) when compared to vehicle-injected Thy1-aSyn mice. MGO treatment of WT littermates had no effect in all behavioural tests performed (Fig. 1A-K, Supplementary Fig. 1D-L).

### MGO aggravates cognitive and olfactory disturbances in Thy1-aSyn mice

Next, we assessed the occurrence of Parkinson’s disease-associated non-motor features in the different animal groups^7,9–11^. The Y maze test was performed to assess spatial reference memory, as a read-out of cognitive function^86,87^. Anxiety-related behaviour was evaluated using the elevated plus maze test^88,89^. Olfactory function was assessed using the block test that evaluates sensitivity to social smells, olfactory acuity, and discrimination^90,91^.

At 20 weeks, Thy1-aSyn mice did not show alterations in the block test (Fig. 1H), Y maze test (Fig. 1I-K), and elevated plus maze (Supplementary Fig. 1K-L) when compared to WT littermates. Likewise, MGO injection did not alter the performance of WT mice in these tests (Fig. 1H-K and Supplementary Fig. 1 F-L). In contrast, MGO-injected Thy1-aSyn mice spent less time sniffing the novel scent (3.2-fold decrease), comparing to vehicle-injected WT mice (Fig. 1H). In addition, MGO-injected Thy1-aSyn mice spent less time (1.4-fold decrease) in the novel arm in the Y maze test (Fig. 1I), comparing to vehicle-injected Thy1-aSyn mice. No differences were observed in anxiety-related behaviour (Supplementary Fig. 1K-L).

Altogether, we observed that MGO exacerbates motor deficits and triggers or anticipates colonic, cognitive, and olfactory disturbances in aSyn-overexpressing mice. We also confirmed that 20 weeks-old Thy1-aSyn mice display impaired motor performance, without colonic, cognitive or olfactory disturbances, compared to WT littermate animals.

### MGO potentiates the accumulation of aSyn in the midbrain, striatum, and prefrontal cortex, and of AGEs in the midbrain of Thy1-aSyn mice

After the behavioural analyses, animals were sacrificed, and the brains were analysed. The left hemisphere was collected and dissected into midbrain, striatum, cerebellum, prefrontal cortex, and hippocampus, and these regions were analysed by immunoblotting. Protein extracts were probed for aSyn and β-actin, for normalization.

Interestingly, MGO-injected Thy1-aSyn mice presented higher levels of aSyn in the midbrain (1.2-fold increase) (Fig. 2A), striatum (1.2-fold increase) (Fig. 2C), and prefrontal cortex (1.2-fold increase) (Fig. 2I) in comparison to vehicle-injected Thy1-aSyn mice.

**Figure 2.**
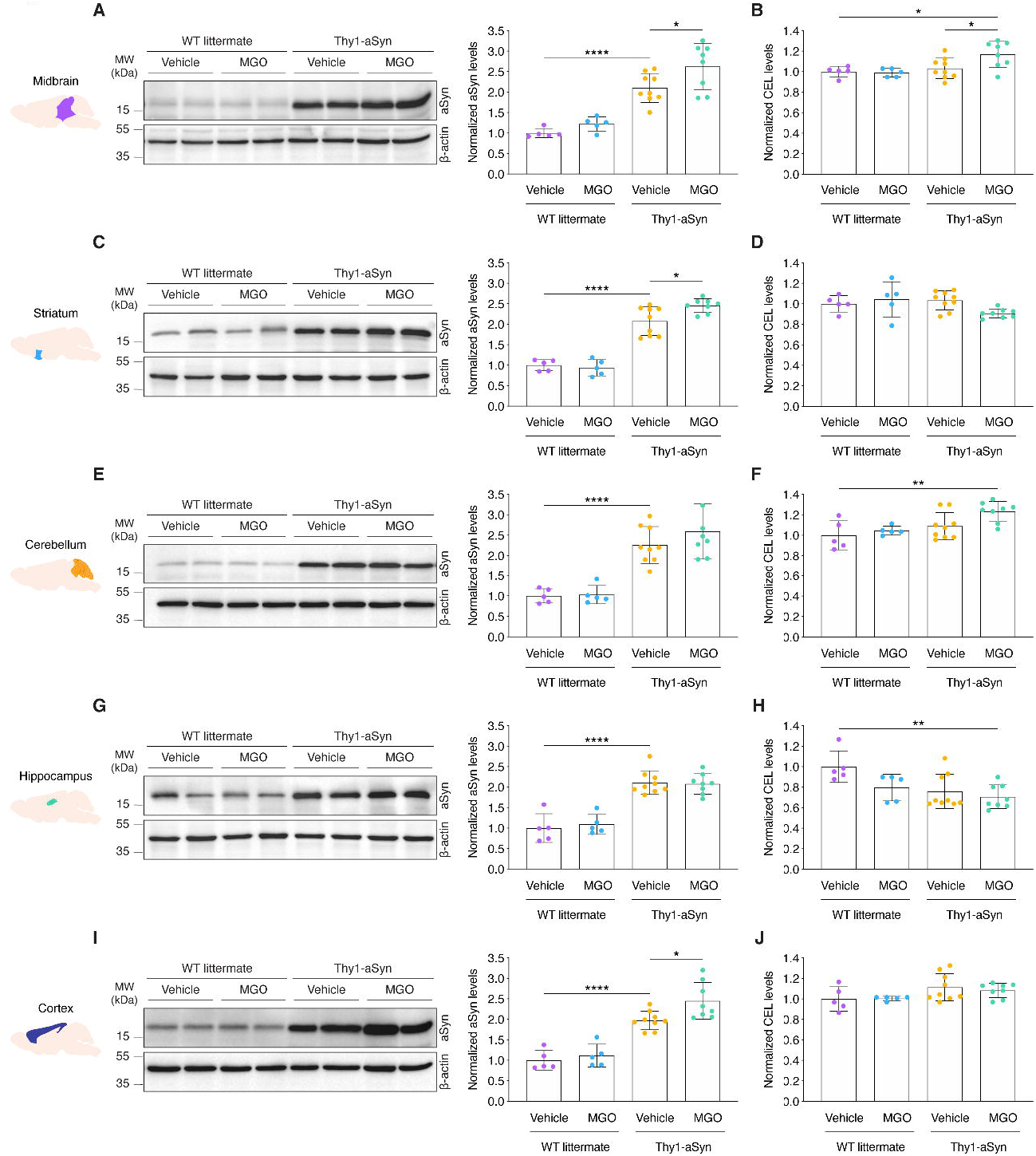
Glycation induces the accumulation of aSyn and AGEs in the midbrain. Wild-type littermate (WT) and Thy1-aSyn transgenic mice received an intracerebroventricular (ICV) injection of MGO or vehicle (PBS) and protein brain extracts from several regions analysed 5 weeks post-injection. Protein extracts were resolved by SDS-PAGE or loaded into membranes in a dotblot system (see Supplemental Figure 2). Membranes were probed with anti-aSyn, anti-CEL and anti-β-actin for normalization. Representative blots showing 2 samples from each experimental group are shown for aSyn, and densitometric analysis represented for aSyn and CEL in **(A, B)** midbrain, **(C, D)** striatum, **(E, F)** cerebellum, **(G, H)** hippocampus, and **(I, J)** prefrontal cortex, respectively. At least n=5 in all groups, Ordinary one-way ANOVA, *p < 0.05, **p < 0.01, ****p < 0.0001.

To understand the possible sustained effects of protein glycation, we measured the levels of CEL, an MGO-derived AGE. MGO-injected Thy1-aSyn mice showed higher levels of CEL, comparing to both vehicle-injected Thy1-aSyn (1.1-fold increase) and to vehicle or MGO-injected WT mice (1.2-fold increase) in the midbrain (Fig. 2B). In the cerebellum, no significant differences were found, except when comparing MGO-injected Thy1-aSyn mice to vehicle-injected WT mice (1.2-fold increase) (Fig. 2F). Strikingly, the hippocampus displayed decreased levels of CEL in both vehicle- and MGO-injected Thy1-aSyn mice (1.3- and 1.4-fold decrease, respectively) in comparison to vehicle-injected WT animals, suggesting that this region might be protected from glycation (Fig. 2H).

As expected, Thy1-aSyn mice presented a 2.1-fold increase in the levels of aSyn in the midbrain, striatum, and hippocampus, and 1.8- and 2.3-fold increase in the prefrontal cortex and cerebellum, respectively, in comparison to WT littermates (Fig. 2). MGO-injected WT mice had no differences in the levels of aSyn or CEL in comparison to vehicle-injected WT mice (Fig. 2).

### MGO triggers neuronal and dopaminergic loss in Thy1-aSyn mice

Next, we compared the number of dopaminergic neurons and of general neuronal population at SNpc surrounding area via immunohistochemical analysis (stereology).

Vehicle-injected Thy1-aSyn mice did not show differences in the population of TH- and NeuN-positive neurons when compared to vehicle-injected WT littermates (Fig. 3A-D). In contrast, MGO-injected Thy1-aSyn mice showed lower number of TH-positive neurons, when compared to vehicle-injected WT littermates (2.1-fold decrease) (Fig. 3B).

**Figure 3.**
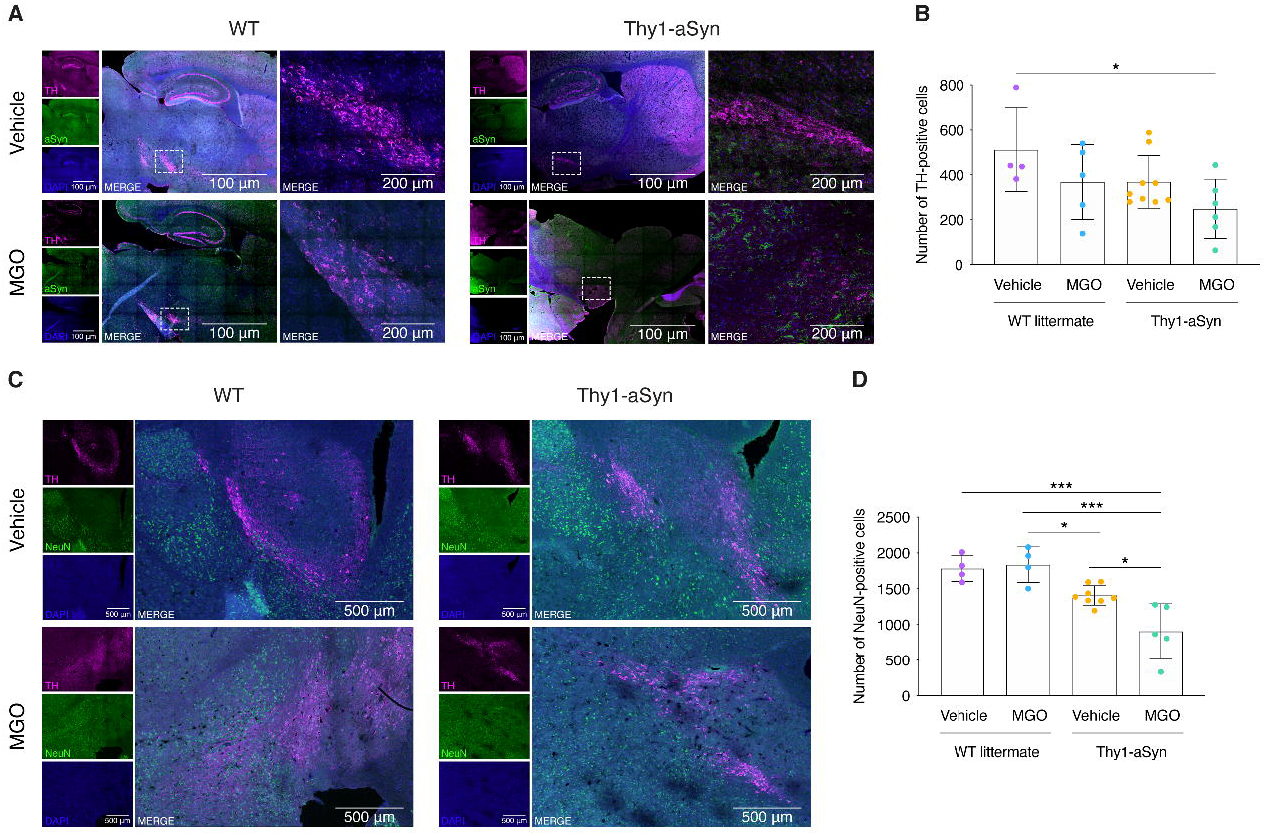
Glycation induces the loss of TH and NeuN neurons in the SNpc. Wild-type littermate (WT) and Thy1-aSyn transgenic mice received an intracerebroventricular (ICV) injection of MGO or vehicle (PBS). **(A)** Representative micrographs of brain sections immunostained for TH (magenta), aSyn (green), and DAPI (blue). Merge signal is shown. Scale bar = 100μm. Depicted area of substantia nigra (dashed square) is shown. Scale bar = 200μm. **(B)** The number of TH-positive cells per experimental group is shown. **(C)** Representative micrographs of brain sections of the substantia nigra immunostained for TH (magenta), NeuN (green), and DAPI (blue). Merge signal is shown. Scale bar = 500μm. **(D)** The number of NeuN-positive cells per experimental group is shown. At least n=4 in all groups, Ordinary one-way ANOVA, *p < 0.05, ***p < 0.001.

addition, MGO-injected Thy1-aSyn mice also had lower number of NeuN-positive neurons in comparison to vehicle-injected Thy1-aSyn mice (1.6-fold) and to both vehicle and MGO-injected WT littermates (2.0-fold decrease) (Fig. 3D).

These findings suggest that MGO treatment results in selective neuronal loss in the SNpc and surrounding midbrain regions of Thy1-aSyn mice.

### Proteomic analysis of differentially regulated proteins

To identify the pathways and molecular mechanisms that are dysregulated upon MGO injection, we performed a proteomic analysis using total protein extracts from the midbrain and prefrontal cortex of 5 animals per group (Thy1-aSyn and WT mice injected with vehicle or MGO). SWATH-based MS was performed, and the MS data matched with a customized library of MS/MS spectra created from LC-ESI-MS analysis of brain protein extracts of animals from each group. Each of the 5 samples per group was analysed 3 times. A total of 2153 proteins were identified for a peptide confidence level of >99%, FDR threshold of 1%. For the quantitative analysis, an initial outlier analysis was performed. The points removed were then replaced with mean values of the analysed group, and iBAQ was normalized. ANOVA analysis between the experimental groups of each brain region was performed to identify the differently regulated proteins.

The response to MGO was followed by comparing MGO-injected with vehicle-injected animals. In the midbrain, 457 proteins were found to be differently regulated between vehicle-injected and MGO-injected WT mice, while 350 proteins were differently present between vehicle-injected and MGO-injected Thy1-aSyn mice. In the prefrontal cortex, 206 proteins were found to be altered by MGO injection in WT mice, and 172 in Thy1-aSyn mice.

The comparison between vehicle-injected Thy1-aSyn and WT mice, revealed the response of the brain proteome to the overexpression of aSyn. Interestingly, we found that 454 proteins were altered in the midbrain, and 256 proteins were altered in the prefrontal cortex of aSyn-overexpressing animals.

#### Proteins uniquely-altered by MGO in aSyn transgenic animals

Next, we identified targets specifically-induced by MGO. To this end, we used a Venn diagram analysis representing the comparisons that enabled us to exclude the general effects of MGO (WT mice injected with MGO vs vehicle) and of aSyn (vehicle-injected Thy1-aSyn vs WT mice), as presented in and for midbrain (Fig. 4A) and prefrontal cortex (Fig. 5A). We found that 160 target proteins were altered as a specific response to MGO treatment in the midbrain, and 105 in the prefrontal cortex of Thy1-aSyn mice. From these, 125 proteins are upregulated, and 35 proteins are downregulated in the midbrain, whereas in the prefrontal cortex 53 proteins are upregulated while 52 are downregulated (Fig. 4B and 5B). By comparing the specific response in the midbrain and prefrontal cortex, we found that only 18 proteins were commonly affected in both regions, suggesting that MGO elicited region-specific alterations.

**Figure 4.**
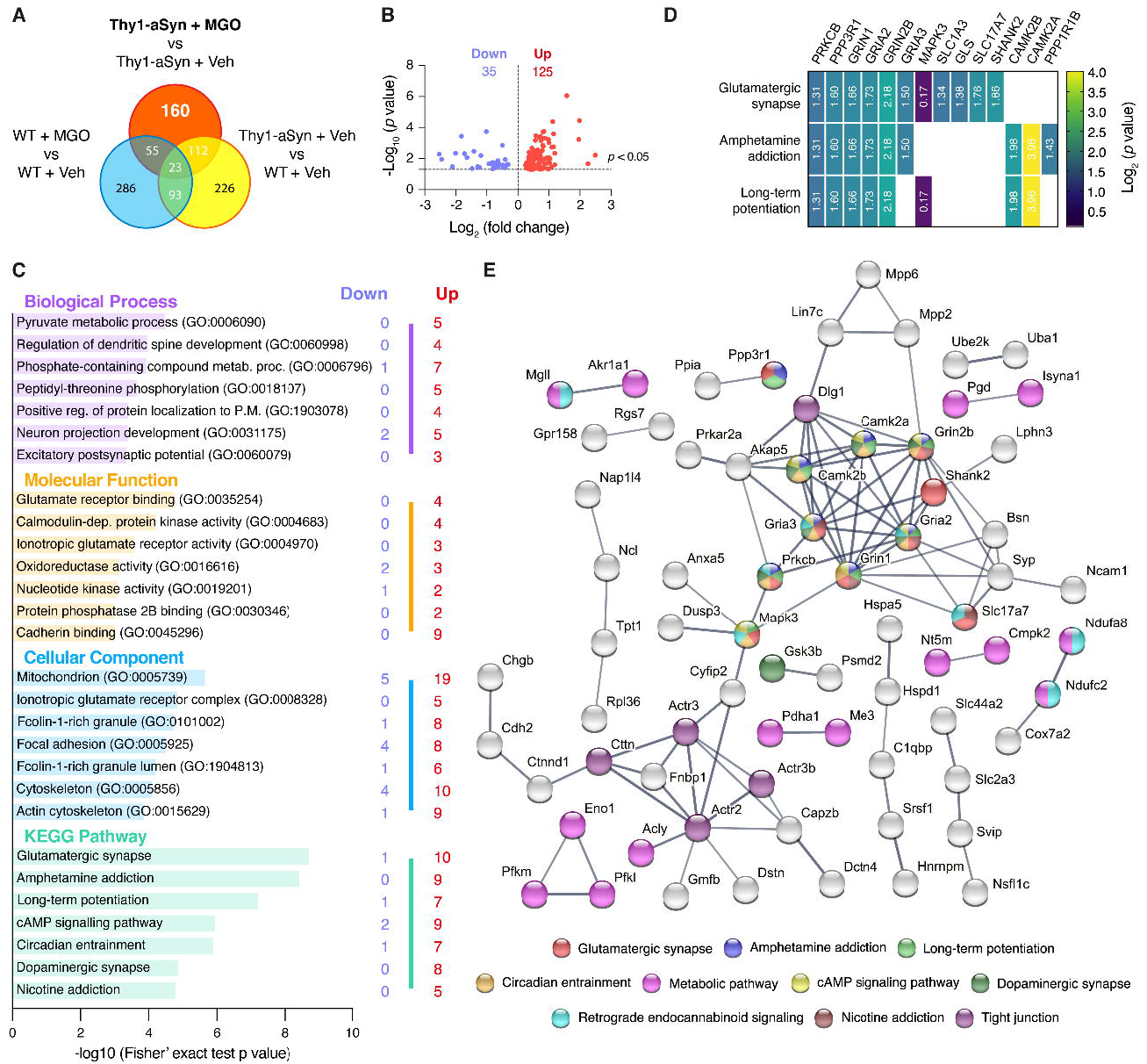
Glycation alters components of the glutamatergic pathway in the midbrain of Thy1-aSyn mice. Proteomic differences between midbrain proteins from WT or Thy1-aSyn mice injected with vehicle or MGO. **(A)** Venn diagram depicting unique and shared midbrain proteome between pairwise comparisons of Thy1-aSyn mice injected with MGO and vehicle, Thy1-aSyn mice and WT mice both injected with vehicle and WT mice injected with MGO and vehicle. **(B)** Volcano plot for unique hits of Thy1-aSyn mice injected with MGO and Vehicle. Statistically significant down and upregulated hits are presented. **(C)** KEEG pathways and gene ontology GO terms analysis is presented, depicting the number of down and upregulated unique hits. Distribution of -log10 (Fisher exact test p value) is shown. **(D)** Heatmap of the uniquely altered proteins corresponding to the top 3 altered KEGG pathways. **(E)** Protein-protein interaction networks of uniquely altered proteins, extracted from the STRING 11.0 database. Only the proteins that are interacting within a network are show. KEGG pathways or GO Biological processes are colour coded.

**Figure 5.**
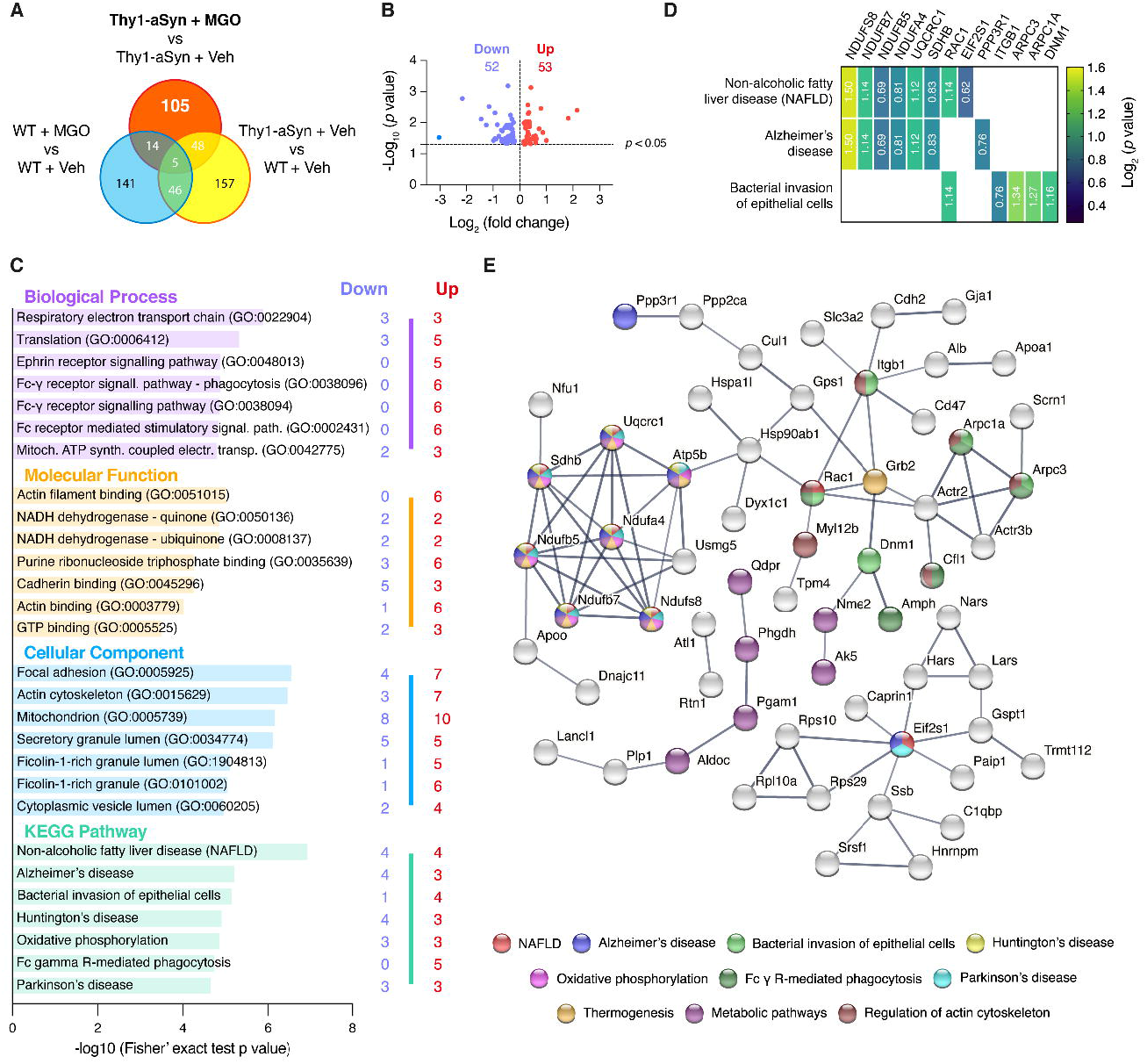
Glycation alters components of the respiratory electron transport chain in the prefrontal cortex of Thy1-aSyn mice. Proteomic differences between cortical proteins from WT or Thy1-aSyn mice injected with vehicle or MGO. **(A)** Venn diagram depicting unique and shared midbrain proteome between pairwise comparisons of Thy1-aSyn mice injected with MGO and vehicle, Thy1-aSyn mice and WT mice both injected with vehicle and WT mice injected with MGO and vehicle. **(B)** Volcano plot for unique hits of Thy1-aSyn mice injected with MGO and Vehicle. Statistically significant down and upregulated hits are presented. **(C)** KEEG pathways and gene ontology GO terms analysis is presented, depicting the number of down and upregulated unique hits. Distribution of -log10 (Fisher exact test p value) is shown. **(D)** Heatmap of the uniquely altered proteins corresponding to the top 3 altered KEGG pathways. **(E)** Protein-protein interaction networks of uniquely altered proteins, extracted from the STRING 11.0 database. Only the proteins that are interacting within a network are show. KEGG pathways or GO Biological processes are colour coded.

#### MGO induces alterations in glutamatergic system-related proteins in the midbrain of Thy1-aSyn mice

Upon functional enrichment analysis, we found that the KEGG pathway mostly affected by MGO in the midbrain of Thy1-mice was the glutamatergic synapse. This was followed by amphetamine addiction, by long-term potentiation. Strikingly, these three pathways share several common proteins (Fig. 4C and D). We found that MGO mainly induced the increase in the levels of proteins related with glutamate production, transport, and its respective transporters and receptors (Fig. 4D).

Consistently with the changes identified in glutamatergic synapses, the most affected GO molecular function was the glutamate receptor binding, and the second most affected GO cellular component was the ionotropic glutamate receptor complex (Fig. 4C). The major changes in GO biological processes were in pyruvate metabolic processes and in the regulation of dendritic spine development (Fig. 4C).

STRING analysis showed a strong connection between the uniquely-altered proteins. A central cluster of highly connected proteins included CAMK2A, CAMK2B, GRIN1, GRIN2B, GRIA2, GRIA3, SHANK2 and PRKCB. Interestingly, other interconnected proteins were associated with tight junction or with metabolic pathway (KEGG) (Fig. 4E).

#### MGO alters components of the respiratory electron transport chain in the prefrontal cortex of Thy1-aSyn mice

In the prefrontal cortex, the most affected KEGG pathways included non-alcoholic fatty liver disease (NAFLD), Alzheimer’s disease, and bacterial invasion of epithelial cells pathways (Fig. 5C and D). Strikingly, the hits related to NAFLD, Alzheimer’s disease, Huntington’s disease, oxidative phosphorylation, and Parkinson’s disease, corresponded to a group of proteins involved in the respiratory electron transport chain (Fig. 5C-E). NADH dehydrogenase [ubiquinone] 1 beta subcomplex subunit 7 (NDUFB7, Ndufb7), NADH dehydrogenase [ubiquinone] iron-sulphur protein 8, mitochondrial (NDUFS8, Ndufs8) and Cytochrome b-c1 complex subunit 1, mitochondrial (UQCRC1, Uqcrc1), were upregulated (Fig. 5D). In contrast, cytochrome c oxidase subunit NDUFA4 (Ndufa4), NADH dehydrogenase [ubiquinone] 1 beta subcomplex subunit 5, mitochondrial (NDUFB5, Ndufb5), and succinate dehydrogenase [ubiquinone] iron-sulphur subunit, mitochondrial (SDHB, SdhB), were downregulated.

Other unique hits and interconnected proteins were associated with actin filament binding, purine ribonucleoside triphosphate binding, cadherin binding processes, Fc γ receptor mediated phagocytosis or to metabolic pathway (Fig. 5C and E).

## Discussion

Synucleinopathies are a group devastating neurodegenerative diseases for which disease-modifying therapies are missing. The misfolding, accumulation, and aggregation of aSyn, a common hallmark among the various synucleinopathies, is thought to be a key event in the neurodegenerative process. However, our poor understanding of the molecular mechanisms underlying neurodegeneration has hampered our ability to develop effective therapies.

The cardinal motor features of Parkinson’s disease result from the loss of dopaminergic neurons in the SNpc, and the consequent depletion of dopamine in the striatum. The degeneration of dopaminergic neurons is believed to be, somehow, associated with the aggregation of aSyn^20–22^. Mutations and polymorphisms have been linked to familiar forms of Parkinson’s disease^92,93^, but account for only 5-10% of cases^94,95^, suggesting that an interplay between genetics and environmental factors account for the vast majority of cases.

Type-2 diabetes mellitus has emerged as an important risk factor for various neurodegenerative disorders and, in particular for Parkinson’s disease^42–44,46,49,50^. Epidemiological studies revealed a strong association between type-2 diabetes mellitus and the risk for developing Parkinson’s disease^44^. Importantly, type-2 diabetes mellitus accelerates the progression of both motor and cognitive deficits in Parkinson’s disease patients^44–46^. However, the molecular mechanisms underlying the connection between the two diseases remains poorly understood. Type-2 diabetes mellitus is a chronic metabolic disease known for glucose metabolism imbalance, and is characterized by hyperglycaemia, insulin resistance, and impaired glucose tolerance^45,47,96^. Remarkably, glycation, a major molecular outcome of type-2 diabetes mellitus, has also been implicated in several neurodegenerative disorders. We found that glycation modulates the pathogenicity of both aSyn and huntingtin, central players in Parkinson’s disease and in Huntington’s disease, respectively^56,97^. Moreover, several studies showed that glycation also plays a role in Alzheimer’s disease^98–100^. Given that protein aggregation is a common hallmark among different neurodegenerative disorders, we posit that glycation may modulate the aggregation and toxicity of the different aggregation-prone proteins and that it may unveil novel targets for therapeutic intervention.

In this study, we investigated how glycation contributes to the dysfunction of neuronal pathways implicated in synucleinopathies, and how it contributes to the onset of Parkinson’s disease-like phenotypes.

To explore our hypothesis, 16 weeks-old Thy1-aSyn or WT littermate mice received a single dose of 160 nmol of MGO or vehicle via ICV injection. In a previous study, we evaluated the impact of an acute delivery of MGO directly to the SNpc or to the striatum^56^. Although several aSyn-associated phenotypes were identified, that experimental procedure was based on the delivery of MGO directly into the brain regions mostly affected in Parkinson’s disease. Here, we applied more physiological amount of MGO (4.5-times lower than previously) and used ICV injection in order to obtain a more homogeneous distribution of MGO in the central nervous system^101–104^. The amount of MGO used corresponds to reported levels of this glycating agent in the mouse brain (approximately 60-130 nmol/brain)^105^. Importantly, the amount used is significantly lower (4 to more than 100 times) than those used in other studies using ICV-delivery ^105–108^.

We performed a detailed behavioural characterization of the animals 4 weeks after ICV injection and, 5 weeks postinjection, animals were sacrificed and brains were collected for biochemical, immunohistochemical and SWATH-MS analysis. The choice of this timeline was based on the reported onset of behavioural alterations in Thy1-aSyn mice^61^.

### MGO injection aggravates Parkinson’s disease like behavioural alterations

Previous reports indicate that most Parkinson’s disease-like behavioural features in Thy1-aSyn mice start at the age of 28 weeks^61^, although they may also exhibit earlier motor deficits^109^. In our experimental conditions, 20 weeks-old Thy1-aSyn mice already presented reduced performance on the vertical pole test, rotarod and wire hang test, and increased hindlimb clasping score^109^. At this stage, no colonic, cognitive, olfactory, and anxiety-related alterations were observed. Therefore, our cohort displayed several alterations that are typically observed at 28 weeks. Moreover, these Thy1-aSyn mice already presented a higher motor impairment in the rotarod test than usual (24 months of age)^110^. In contrast, although Thy1-aSyn mice are reported to be more anxious and to present olfactory alterations at 12 weeks of age, we did not confirm these features in our experimental cohort^90,111^. In agreement with previous reports, no cognitive deficits were observed on the y maze test, as they commonly appear in mice between 28-36 weeks of age^111^.

Following the protocol described, we found that MGO injection in Thy1-aSyn mice aggravates motor deficits as observed in the vertical pole test, rotarod and hindlimb clasping test. Moreover, MGO injection accelerates cognitive impairment in Thy1-aSyn mice, as this was only described to occur between the age of 28 and 36 weeks^111^. In contrast to vehicle-treatment, MGO injection in Thy1-aSyn mice also anticipated the onset of colonic and olfactory disturbances, comparing to vehicle-injected WT animals, assessed in the open field and block test, respectively. These behavioural alterations suggest that MGO accelerates and aggravates disease progression in aSyn-overexpressing mice. In the experimental conditions used, although some tendencies are observed, MGO treatment did not alter the behavioural phenotypes of WT mice. In the future, to test whether glycation triggers Parkinson’s disease-like phenotypes, it will be of interest to design a study to assess the effect of chronic exposure to MGO in WT animals.

### MGO elicits aSyn accumulation and neuronal loss in the midbrain

Our biochemical analysis of different brain regions of interest in the context of Parkinson’s disease showed that MGO-injected Thy1-aSyn mice display increased levels of aSyn in the midbrain, striatum, and prefrontal cortex. This may account for the behavioural alterations observed^112–114^. In agreement with the glycating potential of MGO, we observed an accumulation of AGEs in the midbrain of MGO-injected Thy1-aSyn mice, when comparing to vehicle-injected Thy1-aSyn and to both WT groups.

Chemicals delivered via ICV injection reach various brain regions, including the prefrontal cortex, striatum, hypothalamus, hippocampus, midbrain, and medulla^115^. Although the local concentration of the injected chemical may differ from region to region, the areas surrounding the ventricles should be exposed to higher concentrations of the chemicals, in contrast to subarachnoid areas. Therefore, it was striking to observe that the midbrain was the only region displaying a significant increase of MGO-glycated proteins. Since the striatum and hippocampus should also be exposed to MGO, this finding suggests that either the proteostasis network in the midbrain is less effective in clearing dysfunctional glycated proteins than other brain regions, or that the midbrain is more susceptible to carbonyl-stress. These hypothesis are consistent with the significant decrease in the number of both TH- and NeuN-positive neurons in the SNpc of MGO-injected Thy1-aSyn mice in comparison to vehicle-injected WT mice which, in turn, may account for the observed decrease in motor performance^27,29–31^.

### MGO mainly impacts glutamatergic pathway in the midbrain and oxidative phosphorylation pathway in the prefrontal cortex

Using our proteomics approach, we determined that the response to MGO-injection differs between prefrontal cortex and midbrain. From the uniquely dysregulated proteins specifically induced by glycation in Thy1-aSyn mice, only 18 proteins are commonly dysregulated between the prefrontal cortex and the midbrain. Moreover, the impact of MGO in the midbrain affects a larger number of proteins (350) than in the prefrontal cortex (172), for the same total amount of detected proteins. Although MGO increased the levels of aSyn in both regions, the differences may be due to a distinct exposure to MGO, as AGEs accumulated in the midbrain, but not in the prefrontal cortex. Another possibility is that this may also be the result of a higher susceptibility of midbrain cells to MGO, as previously suggested.

MGO glycation is known to trigger the unfolded protein response, to impair oxidative metabolism, to drive mitochondrial dysfunction, and to impact on the oxidative stress response^49,50,116,117^. Several of these consequences are intrinsically related with the depletion of cellular defences against oxidative stress since reduced glutathione and NADPH are shared cofactors to cope with both oxidative stress response and with the detoxification of MGO by glyoxalases and aldose reductases^49,50^. This is particularly relevant in neurons, which are more vulnerable to MGO due to their lower capacity to detoxify this compound, particularly when compared to astrocytes^118,119^. Failure in its detoxification results in increased glycation of several proteins, particularly in the mitochondria^120^. In fact, neuronal mitochondrial damage is well established, and is known to suppress oxygen consumption, decrease the activity of respiratory chain complexes, and the capacity of energy production, increasing the production of reactive oxygen species (ROS)^121,122^. Importantly, mitochondrial dysfunction in the dopaminergic neurons is associated with Parkinson’s disease^123,124^. Thy1-aSyn mice injected with MGO displayed severe dysregulation of oxidative phosphorylation in the prefrontal cortex. Several elements of mitochondrial complex I were dysregulated, with a decrease of NDUFB5 and increase of both NDUFB7 and NDUFS8. Moreover, components of complex II (SDHB) and complex IV (NDUFA4) were decreased, while a component of complex III (UQCRC1) was increased. These findings suggest that increased levels of MGO in the brain may contribute to mitochondrial dysfunction in the cortical area.

Altered glutamatergic firing has been described in Parkinson’s disease and is believed to be caused by the dopamine depletion in the striatum^27–31^. The SNpc is highly rich in NMDA, AMPA and metabotropic glutamate receptors, and glutamate or glutamate-dopamine neurons are intermixed with midbrain dopaminergic neurons^34–36^. In the midbrain of MGO-injected Thy1-aSyn mice, we observed a generalized loss of neurons, and a specific loss of dopaminergic neurons. Moreover, we found that several uniquely dysregulated proteins in the midbrain of MGO-injected mice belong to the glutamatergic pathway, which may reflect a possible increase of glutamatergic signalling. Specifically, there was an increase of the levels of glutaminase and VGLUT1. These events suggest a probable increase in the presynaptic production of glutamate, and of its storage in glutamatergic vesicles (Fig. 6). The levels of astrocytic EAAT1 were also increased and this protein is classically responsible for the rapid removal of released glutamate from the synaptic cleft^125,126^. This finding further supports the hypothesis that glutamate production and release are increased in this brain region upon MGO insult. Moreover, a generalized increase of AMPA (GRIA2 and GRIA3) and NMDA (GRIN1 and GRIN2B) receptors was observed, most probably at the post-synaptic neuron (Fig. 6B and C). In particular, we observed an increase of the GRIN2B, a subunit of NMDA receptor that is generally linked to cell death signalling^127,128^, and is typically more abundant early in development, shifting to GRIN2A during development^128^. Our study does not allow us to resolve whether the increased levels of these glutamate receptor subunits are at the post-synaptic membrane (Fig. 6B), extrasynaptically (Fig. 6C), or at the cytosol. Nevertheless, several post-synaptic glutamatergic targets are also increased, including the calcineurin subunit B type 1, PPP1R1B, PKCβ, and SHANK2, suggesting increased glutamatergic activity. In fact, we also found an increase in the levels of CAMK2A and CAMK2B in MGO-injected Thy1-aSyn mice, which are known to promote the trafficking and transient translocation of AMPA receptors to the postsynaptic membrane^129^. Together with the observation of decreased levels of MAPK3/ERK2, these findings suggest an impairment of LTP^130^, as we previously reported^129^. Excessive release of glutamate may also activate extrasynaptic glutamate receptors. Oligomeric aSyn, may not only induce increased release of glutamate from astrocytes, but also activate extrasynaptic NMDA receptors, and this process may lead to synaptic loss. Importantly, the specific extrasynaptic NMDA receptor antagonist NitroSynapsin protects from these deleterious effects of oligomerized aSyn^131^. Independently of the localization of glutamate receptors, excessive glutamate may trigger apoptotic cell death because of intracellular calcium overload upon receptor stimulation, and is a common hallmark of neurodegenerative diseases^132^.

**Figure 6.**
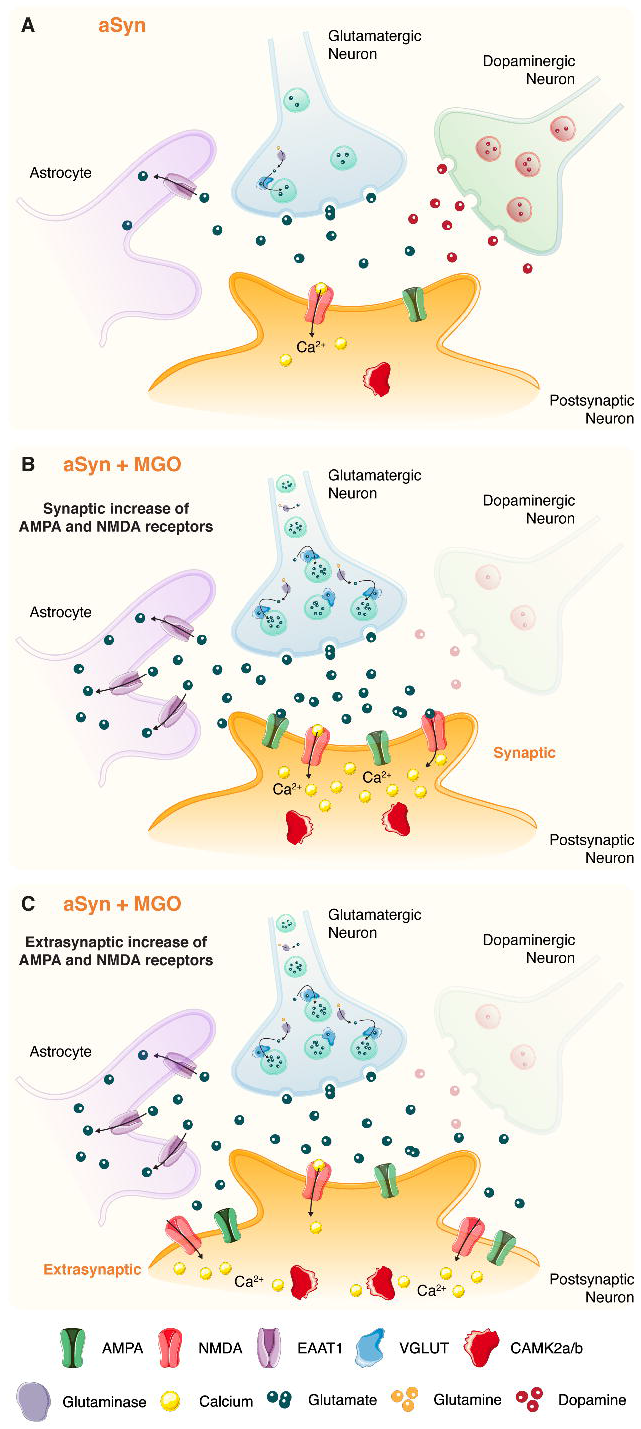
Effects of MGO in the midbrain of Thy1-aSyn mice. The diagram depicts the MGO-altered glutamatergic proteins in the midbrain of Thy1-aSyn mice. Schematic of dopaminergic and glutamatergic signalling in **(A)** vehicle-injected mice, **(B)** and **(C)** MGO-injected mice. While in **(A)** vehicle-injected mice, basal dopaminergic and glutamatergic signalling should occur, **(B)** and **(C)** MGO-injected mice present a smaller number of dopaminergic synapses. **(B)** and **(C)** Proteomics data suggest a probable increase of glutamate production (glutaminase) and packaging in glutamatergic vesicles (VGLUT), the release of glutamate to the synaptic cleft and astrocytic reuptake (EAAT1) or entry to the postsynaptic neuron via AMPA or NMDA receptors. The increase of these receptors may occur at the **(B)** synapse or **(C)** extrasynaptically, inducing calcium entry to the cell. An increase of both CAMK2a and CAMK2b is also observed, supporting the hypothesis of enhanced glutamatergic signalling. A strong imbalance in the post-synaptic levels of calcium can occur and contribute to excitotoxic events.

Based on our findings, we cannot distinguish whether the alteration in glutamatergic signalling precedes dopaminergic degeneration or is a consequence of dopaminergic dysregulation in MGO-injected Thy1-aSyn mice. In a first scenario, MGO could exert a direct effect that elicits glutamatergic hyperactivity, causing an excitotoxic phenomenon that triggers general neuronal degeneration and specific dopaminergic loss. Dopamine depletion would explain the cardinal features of Parkinson’s disease, and are consistent with the motor alterations we observed. In agreement with this hypothesis, we observed the accumulation of aSyn in the midbrain, and we previously reported that MGO-treatment increases both aSyn glycation and oligomerization^56^. Moreover, we further reported that prolonged exposure to aSyn oligomers drives to the long-lasting increase of basal glutamatergic synaptic transmission^129^. In fact, aSyn oligomers may activate NMDA receptors, increasing the basal levels of intracellular calcium and recruiting AMPA receptors to the membrane^129^. In a second scenario, MGO could first trigger the loss of dopaminergic neurons, possibly due to the enhanced accumulation aSyn. In an indirect process, the brain may respond to dopamine depletion by increasing glutamatergic firing in the midbrain. This compensatory mechanism should enhance the release of dopamine from the surviving dopaminergic neurons to maintain dopamine homeostasis. It is presently unclear if the glutamatergic alterations cause or exacerbate neurodegeneration in patients with Parkinson’s disease. However, glutamatergic hyperactivity may prompt an excitotoxic cascade further promoting neurodegeneration^37,132^, which would ultimately lead to the observed behavioural alterations. Importantly, alterations in glutamate content in the brain of Parkinson’s disease patients have been reported by several clinical studies using magnetic resonance imaging (MRI), positron emission tomography (PET) and single photon emission computed tomography (SPECT), consistent with increased glutamate neurotransmission^133–135^. Moreover, increased levels of glutamate in the plasma of Parkinson’s disease patients have been reported, further reflecting increased cerebral glutamatergic activity^136,137^. Alterations in this glutamatergic transmission contribute to the pathophysiology of dyskinesias, impaired motor coordination and motor fluctuations^28,138–142^. Additionally, dysfunction of glutamate metabolism is also implicated in non-motor features of Parkinson’s disease, including depression and cognitive impairment^143–145^.

## Conclusions

The pathogenesis of Parkinson’s disease and the mechanisms leading to the accumulation and aggregation aSyn remain poorly understood. Type-2 diabetes mellitus is an important risk factor for Parkinson’s disease and, interestingly, glycation mediates aSyn pathogenesis in vitro and in animal models. In this study, we demonstrate that glycation exacerbates Parkinson’s disease-like motor and non-motor features in Thy1-aSyn mice. Importantly, these behavioural alterations are followed by the accumulation of aSyn in the midbrain, striatum, and prefrontal cortex, and by neuronal loss in the midbrain. Furthermore, we found that MGO-injected Thy1-aSyn mice show pronounced alterations in proteins of the glutamatergic pathway within the midbrain, consistent with increased production of glutamate and increased glutamatergic transmission. Thus, we suggest that in conditions of aSyn pathology, MGO induces glutamatergic hyperactivity in the midbrain directly, or indirectly by depleting dopaminergic neurons, which aggravates both motor and non-motor features in mice^40,138^.

In conclusion, we uncovered a major role for MGO-derived glycation in the exacerbation or anticipation of Parkinson’s disease-like features, suggesting that anti-diabetic/anti-glycation agents hold promise as disease-modifying agents in Parkinson’s disease. Likewise, glutamatergic-silencing molecules may suppress excitotoxic events that underlie neuronal degeneration in synucleinopathies.

## Abbreviations

aSyn: Alpha-synuclein
AGEs: advanced glycation end products
AMPA: α-amino-3-hydroxy-5-methyl-4-isoxazolepropionic acid
BG: basal ganglia
CEL: N(epsilon)-(carboxyethyl)lysine
EAAT1: excitatory amino acid transporter
GO: Gene Ontology
ICV: intracerebroventricular
KEGG: Kyoto Encyclopaedia of Genes and Genomes
LB: Lewy bodies
LN: Lewy neurites
MGO: methylglyoxal
NMDA: N-Methyl-D-Aspartate (NMDA)
PTMs: posttranslational modifications
SNpc: substantia nigra pars compacta
SWATH-MS: sequential window acquisition of all theoretical mass spectra
TH: tyrosine hydroxylase
VGLUT: vesicle glutamate transporter

## Author contributions

AC, JEC, PGA, LVL, TFO, and HVM designed research; AC, JEC, MG, MTF, IMM, BFG, BMA, RAG, LS performed research; PRF prepared and provided materials; BMA, RGA and RM made statistical analysis of proteomic data; AC, BMA, RAG and HVM performed proteomic analysis; AC and HVM analysed all data; AC and HVM wrote the manuscript. All authors reviewed the manuscript.

## Acknowledgements

We would like to acknowledge the vivarium and behavioural facilities at Instituto de Medicina Molecular – João Lobo Antunes for all the support. We also acknowledge Prof. Rosalina Fonseca, Prof. Sílvia V. Conde, Prof. Rita Machado de Oliveira, Dr. Natália Madeira, Dr. Liliana Lopes, Dr. Tatiana Burrinha, and Dr. Catarina Perdigão for fruitful discussions. We acknowledge Dr. Manuela Correia for all the laboratory support.

## Funding

This study was supported by Fundação para a Ciência e Tecnologia (FCT) PTDC/NEU-OSD/5644/2014, iNOVA4Health UID/Multi/04462/2013 and Sociedade Portuguesa de Diabetologia. Authors were supported by: AC (FCT, PD/BD/136863/2018; ProRegeM – PhD programme, mechanisms of disease and regenerative medicine); LS (SFRH/BD/143286/2019; PhD Studentship); BFG (PTDC/NEU-OSD/5644/2014). TFO is supported by the Deutsche Forschungsgemeinschaft (DFG, German Research Foundation) under Germany’s Excellence Strategy - EXC 2067/1- 390729940, and by SFB1286 (B8).

## Competing interests

The authors have no conflicts of interest to declare.

## Supplementary material

Supplementary material includes Extended Material and Methods and Supplementary Figures 1-4.

---

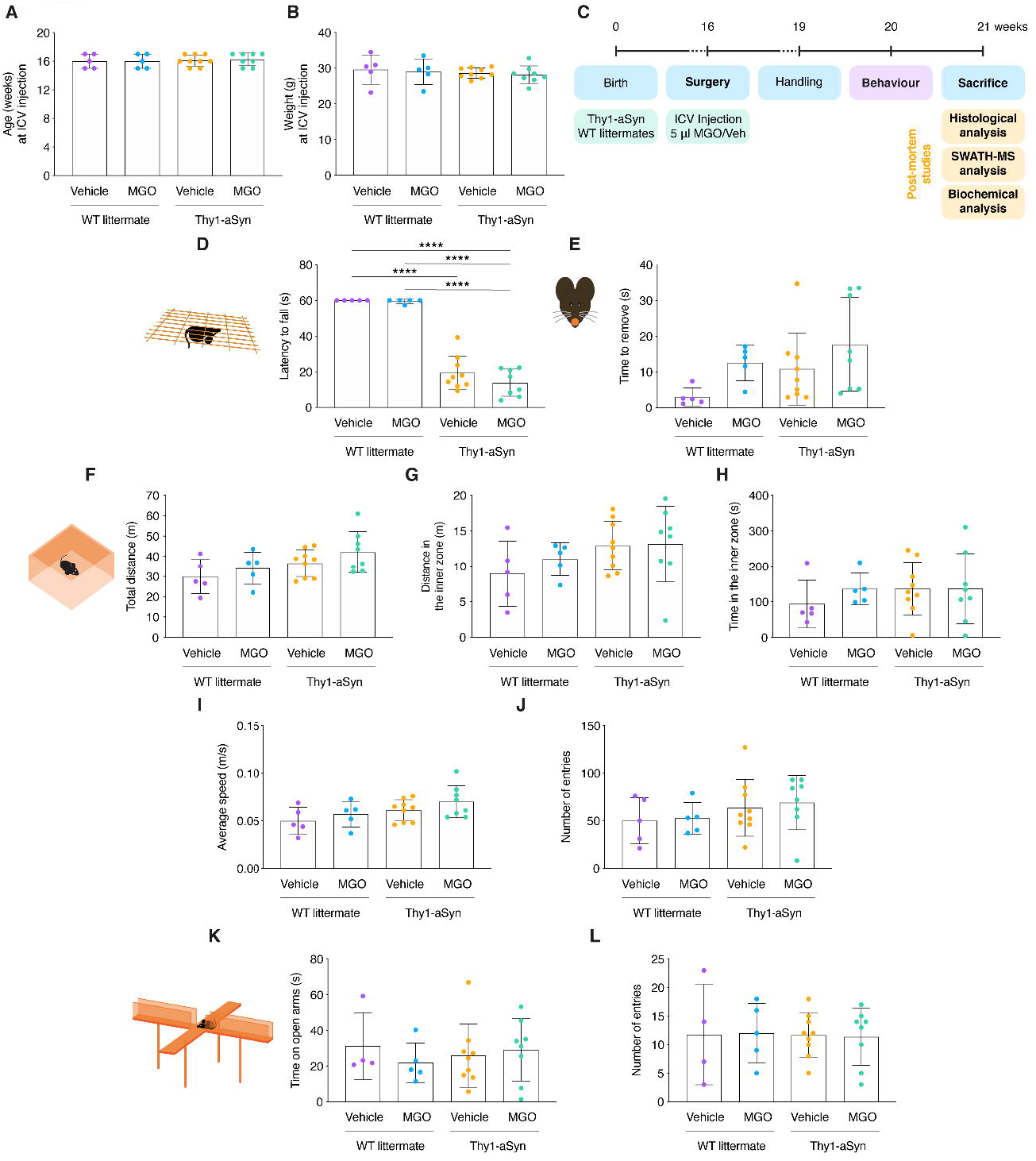

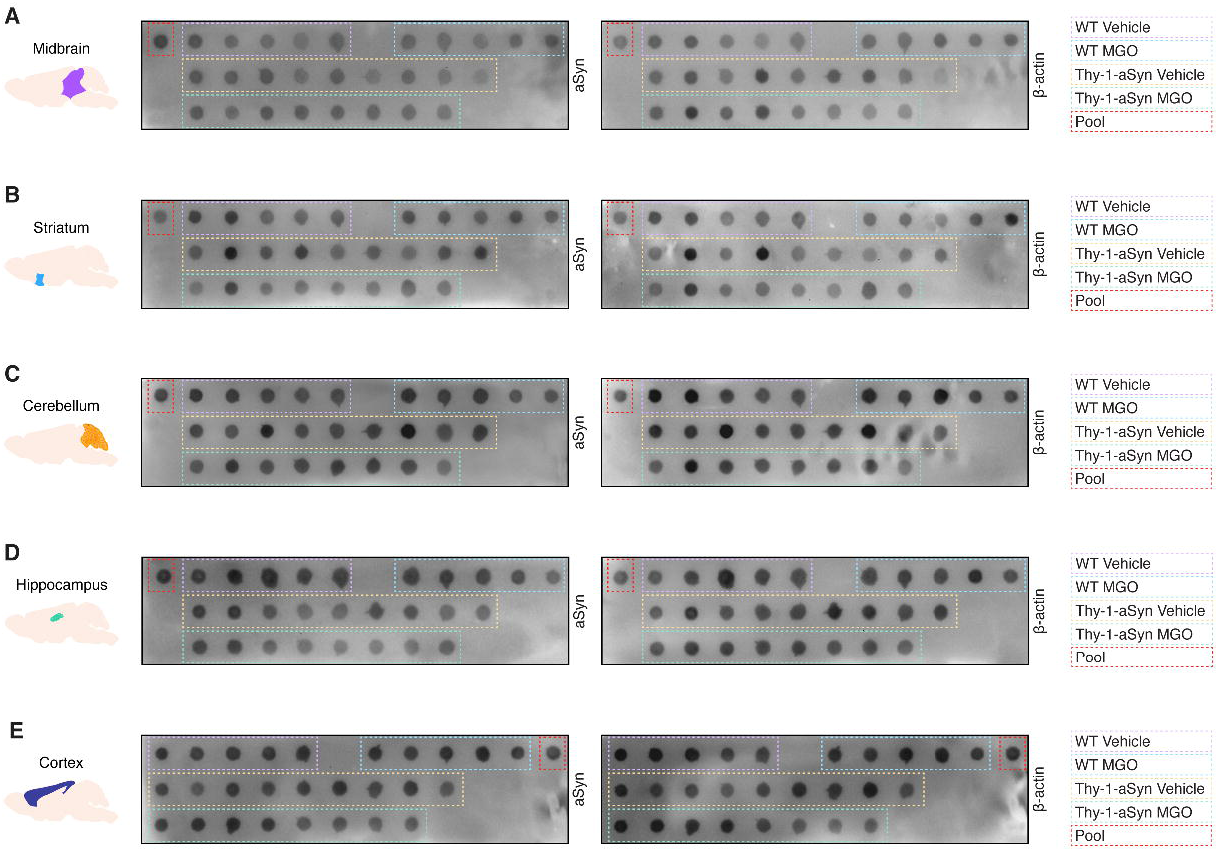

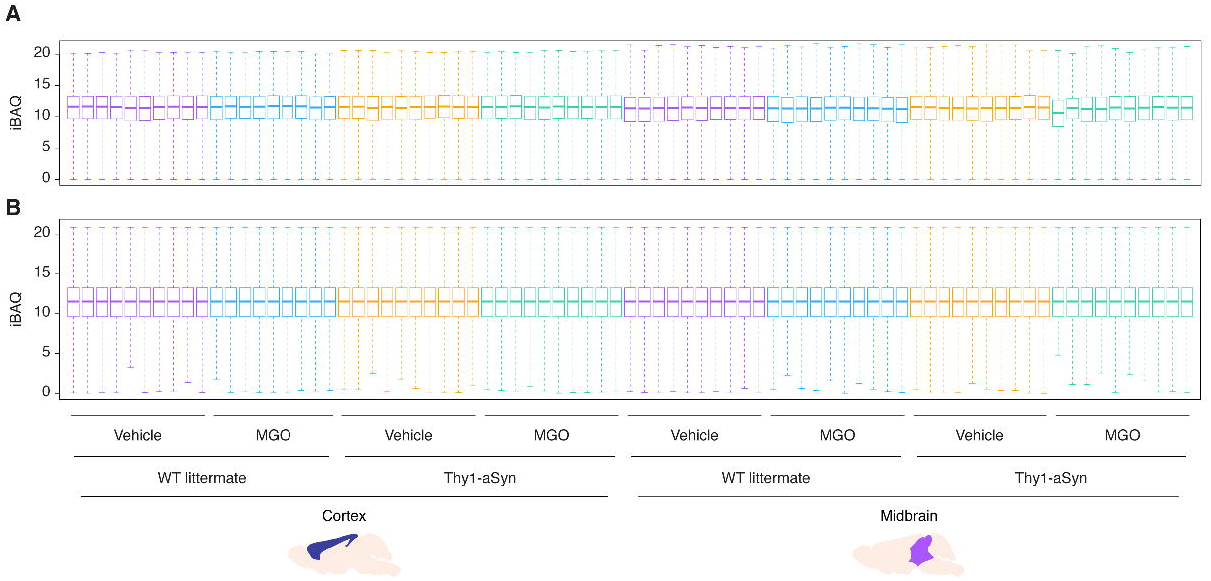

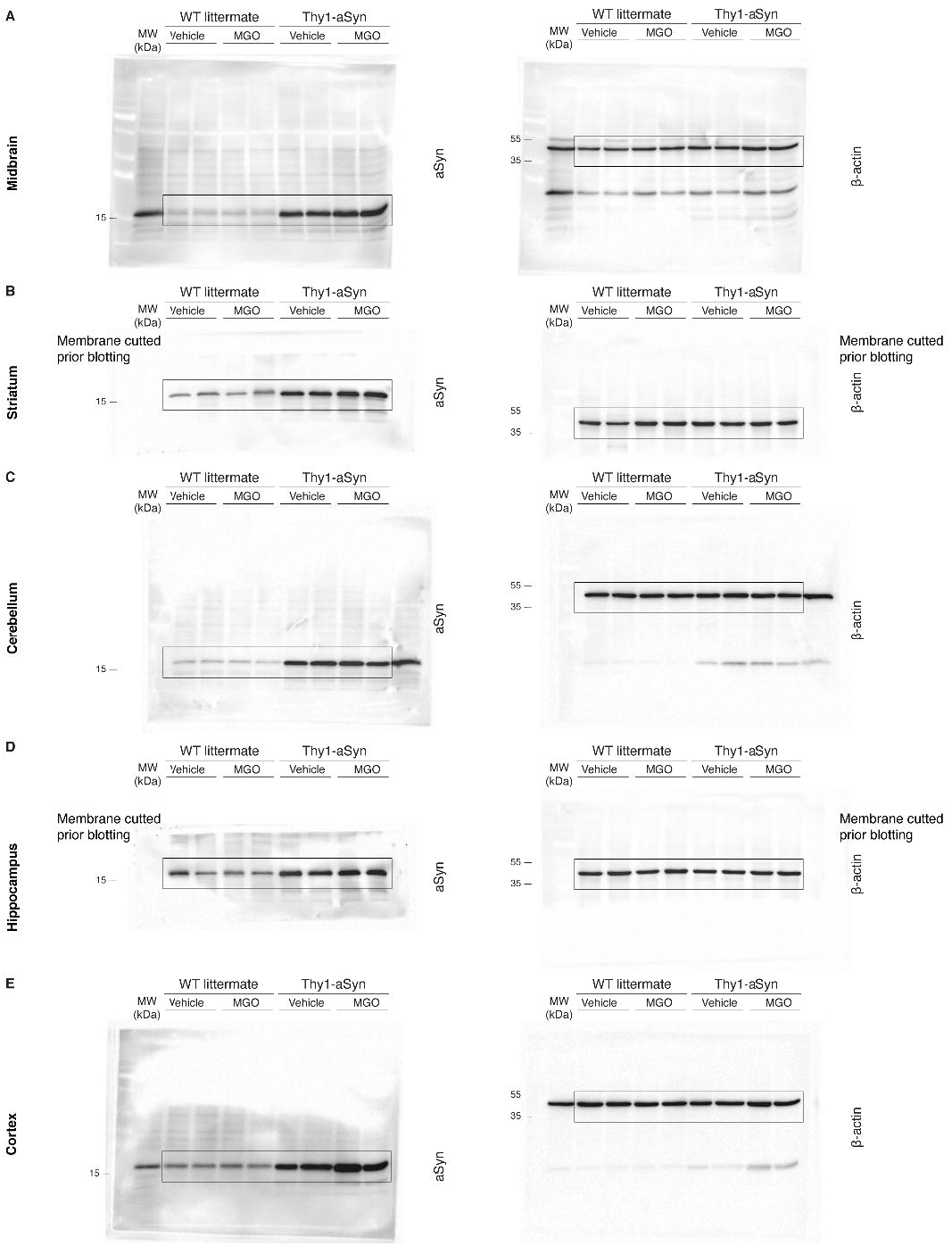

## Notes

### Competing Interest Statement

The authors have declared no competing interest.

