## Supplementary material for "Glycation modulates glutamatergic signalling and exacerbates Parkinson’s disease-like phenotypes"

#### Extended Material and Methods

##### Methylglyoxal production and standardization

Purified MGO was diluted to estimated 50-100 µM in a solution with 1 mM aminoguanidine hydrochloride in 50 mM sodium phosphate buffer (pH 7.4). The mixture was incubated at 37ºC for 4 hours. The reaction between MGO and aminoguanidine forms aminotriazine, whose absorbance was measured spectrophotometrically at 320 nm, from which the concentration of MGO is deduced, according to $\varepsilon_{320}$ = 2411 $M^{-1}.cm^{-1}$.

##### Animals

Animal procedures were carried out in accordance with the European Community guidelines (Directive 2010/63/EU), Portuguese law on animal care (DL 113/2013), and approved by the iMM Internal Committee and the Portuguese Animal Ethics Committee (Direcção Geral de Alimentação e Veterinária - DGAV).

Animals were maintained under controlled light (12 hours light/12 hours dark cycle) and environmental conditions, with a constant temperature of 21 ± 0.5 ºC, and relative humidity of 60 ± 10%, and had free access to commercial chow and water (ad libitum). Mice were housed in groups, with 2 to 5 animals per cage. Only male animals were used in all experimental procedures. Mice were sacrificed by exsanguination, through the perfusion of PBS from the heart, after anaesthesia under isoflurane atmosphere.

Transgenic mice overexpressing human aSyn under the Thy-1 promoter were generated on a mixed C57BL/6-DBA/2 background as described previously[1]. Animals were obtained from our breeding colony on this background by breeding mutant females with wildtype (WT) C57BL/6-DBA/2 males. Offspring were genotyped via polymerase chain reaction (PCR) amplification analysis of DNA extracted from ear or toe. PCR was performed using the following primers: Thy-1- F: 5′-CTG GAA GAT ATG CCT GTG GA-3′, Thy-1-R: 5′-GAG GAA GGA CCT CGA GGA AT-3′, with an annealing temperature of 60ºC and 40 cycles of amplification as previously[2].

##### Intracerebroventricular injection of MGO and general procedures

The bolus injection was performed using a Hamilton syringe, attached to a micropump system with a flow rate of 0.5 µL/min. Following the surgery, animals were allowed to recover from the procedure. Three weeks post-injection, mice were weighed, handled, and general phenotype assessed by SHIRPA analysis. Behavioural testing was performed starting four weeks after surgery. Upon conclusion of the behavioural phenotyping, mice were sacrificed (Supplementary Fig. 1C). Brains were collected and separated into left and right hemispheres: the right hemisphere was transferred into paraformaldehyde (PFA) solution for fixation and immunohistological analysis; the left hemisphere was dissected and rapidly frozen in liquid nitrogen for biochemical and sequential window acquisition of all theoretical mass spectrometry (SWATH-MS) analysis. Baseline body weight prior to the surgery was measured. Body weight was recorded immediately before mice sacrifice.

##### Behavioural tests

Mice were handled once a day for 5 days prior to behaviour evaluation. Mazes were cleaned with a 10% ethanol solution between each animal. All behavioural tests were done during the light phase between 8 a.m. and 9 p.m.

in a sound attenuated room. Prior to each test, animals were allowed to acclimate to the room for 30 minutes.

##### SHIRPA protocol

SHIRPA protocol was used to evaluate general health behaviour and phenotype characterization. We evaluated hindlimbs clasping and colonic function. Hindlimbs clasping is a marker of disease progression and cerebello-cortico-reticular and cortico-striato-pallido-reticular pathways function assessment and was performed as previously described[3]. In this test, the mouse was suspended by the tail and the extent of hindlimb clasping observed for 30 seconds and scored from 0 to 3: score 0 – both hindlimbs were splayed outward away from the abdomen with splayed toes; score 1 – one hindlimb was retracted or both hindlimbs were partially retracted; score 2 – both hindlimbs were partially retracted toward the abdomen and were touching the abdomen; score 3 – both hindlimbs were fully clasped and touching the abdomen.

Colonic function was assessed during open field test[3,4]. The number of faecal pellets were counted after 10 minutes in the open field arena.

##### Open field test

The open field test was used to observe general motor activity, gross locomotor activity, and exploration habits[5]. Assessment took place in a square arena (40 cm length x 40 cm wide x 40 cm height) with opaque walls, and a single trial was done. The mouse was placed in the centre of the arena and allowed to freely explore for 10 minutes, while being recorded by an overhead camera. The footage was then analysed by an automated tracking system and distance moved, velocity, and time spent in pre-defined zones were measured. By observation, we recorded rearing (standing up on hind limbs) and grooming behaviours, and defecation and urination.

##### Pole test

The pole test was performed to assess locomotor activity, mainly motor coordination, and balance[3,5-7]. The pole was composed of metal rod with 50 cm length and a diameter of 1 mm wrapped with paper tape. The base of the pole was placed in a cage filled with bedding material. The mouse was placed head-upward close to the top of a vertical pole and was expected to orient downward and descend the length of the pole back into the cage. A maximum time of 180 seconds was given to complete the task. Each mouse underwent one training and four trials, with 30 minutes intervals. Training and trial tests were recorded. The time to turn down, to climb down and total time were measured manually. Data from four trials were averaged for each mouse and presented as average plus standard deviation.

##### Rotarod test

The rotarod test was used to evaluate motor coordination, balance, and motor learning[3,5-7]. A commercial apparatus with a rat rod with a diameter of 6 cm was used. Animals underwent one training and three trials at 30 minutes intervals. During the training, the mouse was placed on the rotating rod at 7 rpm until it could stand on the rod (about 2–3 minutes). For testing, the mouse was placed on a rotating rod with either constant rotation (7 rpm) or continuous acceleration (from 4 to 40 rpm in 5 minutes). In the steady rotation protocol, the animals were placed on the rotating rod at a constant speed of 7 rpm, for three minutes. In accelerating conditions, the mouse was placed on the rotating rod at a constant speed of 4 rpm that gradually accelerates from 4–40 rpm in 10 minutes. The latency to fall was recorded for both protocols. Data is presented as average of the three trials for each mouse.

##### Wire hang test

Wire hang test was used to evaluate balance and grip strength[5,8]. The mouse was placed hanging from an elevated wire cage top, which was then inverted and suspended above the home cage (1 m) for 60 seconds. Animals underwent three trials at 30 minutes intervals. The latency to fall was recorded. Data is presented as average of the three trials for each mouse.

##### Adhesive removal test

Adhesive removal test was performed to evaluate sensory and motor deficits related to the paw and the mouth[9]. This test consists of applying a white adhesive tape of 0.8 mm of diameter onto the snout of the mouse. The animal is then released and the time-to-remove the adhesive was measured. Each mouse underwent one training and three trials with 15 minutes intervals. Data is presented as average of the three trials for each mouse.

##### Y maze test

Y maze test was used to assess short-term spatial reference memory, which is hippocampal dependent[10,11]. The test was performed in a Y-shaped maze with three arms (arm A - 20 cm length x 5 cm wide x 12 cm height, arms B and C - 15 cm length x 5 cm wide x 12 cm height), angled at 120º and with opaque walls. During the training, the mouse was placed in the start point of the Y maze with a closed arm and allowed to freely explore it for 6 minutes. After one hour, the "novel" arm was open, and the mouse allowed to freely explore the maze for 5 minutes. A single trial was done and recorded by an overhead camera. Time spent in the novel arm was measured manually, as well as the number of triads (i.e., ABC, CAB, or BCA but not ABB) and entries. The alternative behaviour score (%) for each mouse was calculated as the ratio of the number of alternations to the possible number (total number of arm entries minus two) multiplied by 100. The maze was cleaned with diluted 10% ethanol between tests to eliminate odours and residues. Data presented result from a single trial for each mouse. One animal from vehicle-treated WT littermate group was considered an outlier since it deviated more than 1 standard deviation from the group mean of time spent on novel arm. One mouse from MGO-treated Thy1-aSyn group failed to complete the test since it was too stressed.

##### Elevated plus maze test

To characterize anxiety-related behaviour, the elevated plus maze test was used[12,13]. The apparatus has a small central platform with four arms radiating outwards and is raised above the ground to a height of 75 cm. The arms are placed at an angle of 90º from each other and are 50 cm in length and 10 in width. Alternating arms are enclosed by high opaque walls of 50 cm height, with open tops. The mouse was placed in the centre of the maze and allowed to freely explore for 5 minutes for a single trial and recorded by an overhead camera. The time spent on open arms and the number of entries were measured manually. Data presented result from a single trial for each mouse.

##### Block test

We performed the block test to assess olfactory function[14,15]. This test evaluates sensitivity to social smells, an ethologically essential ability in mice, thus measuring the olfactory acuity and discrimination. Housed animals were exposed to five wood blocks (2 cm) placed inside each cage for 7 days. During this period, the cage bedding was not replaced. Upon testing, four blocks originally from the mouse's own cage were placed into a new cage, approximately 1-2 cm apart. The mouse was placed on the novel cage and videotaped for 30 seconds. Each mouse underwent four training sessions. On the trial test, one block was replaced by a block that was originally in a cage with a different set of animals. The mouse was videotaped for 1 minute. The time sniffing novel scent was measured manually. Data presented result from a single test trial for each mouse. Two mice from vehicle-treated Thy1-aSyn group displayed freezing behaviour and failed to perform the test.

#### Tissue lysate preparation

200 µl of RIPA buffer (50 mM Tris-HCl pH 7.4, 150 mM NaCl, 2 mM EDTA, 0.1 % SDS, 0.25 % sodium deoxycholate) were added per 0,02 g of brain tissue. Samples were macerated with an automatic pestle. Samples underwent three cycles of sonication in pulses (1 second on, 45 milliseconds off, for 30 seconds, with 15% of intensity), with 1 minute incubation on ice between them. Protein extracts were centrifuged for 10 minutes at 9.600 rcf at 4 ºC to pellet tissue and cell debris. Supernatant was collected and total protein was quantified using Pierce® BCA Protein Assay Kit (Thermo Fisher Scientific; Waltham, MA, USA).

#### SWATH-MS analysis

A high-throughput proteomics analysis using NanoLC coupled to the TripleTOF 6600 (at UniMS, Mass Spectrometry Unit at iBET/ITQB) was performed to screen for differences in protein expression between experimental groups of midbrain or frontal cortex protein extracts.

##### Sample preparation for mass spectrometry analyses

Eighty micrograms of each sample were reduced with 10 mM dithiothreitol (BioUltra, Sigma) for 45 min at 56 °C, alkylated with 20 mM iodoacetamide (BioUltra, Sigma) for 30 min at room temperature in the dark, and then precipitated with acetone (HPLC Plus, Sigma) overnight at -20 °C. The dried sample was resuspended in 50 mM ammonium bicarbonate (BioUltra, Sigma) and digested overnight at 37 °C with trypsin (Sequencing Grade Modified Trypsin, Promega) in a 1:50 trypsin:sample ratio. A second digestion was performed for 3 h at 37 °C, by adding trypsin in a 1:100 trypsin:sample ratio with 80% (v/v) acetonitrile (Optima LC/MS grade, Fisher Scientific). The sample was dried on a SpeedVac (ThermoSavant Scientific) and resuspended in 5 % Formic acid (Optima LC/MS grade, Fisher Scientific) to perform peptide cleanup using C18 microcolumns (OMIX C18 pipette tips, Agilent), and then dried again.

##### Information-dependent acquisition (IDA) runs to generate the spectral library

Nano-liquid chromatography-tandem mass spectrometry (nanoLC-MS/MS) analysis was performed on an ekspert™ NanoLC 425 cHiPLC system coupled with a TripleTOF 6600 with a NanoSpray III source (Sciex Framingham, US). Peptides were sprayed into the MS through an uncoated fused-silica PicoTip™ emitter (360 µm O.D., 20 µm I.D., 10 ± 1.0 µm tip I.D., New Objective, Oullins, France). The source parameters were set as follows: 15 GS1, 0 GS2, 30 CUR, 2.5 keV ISVF, and 100 $^{0}$C IHT. Peptides were separated through reversed-phase chromatography (RP-LC) in a trap-and-elute mode. Trapping was performed at 2 µL/min on a NanoLC Trap column (Eksigent 350 µm x 0.5 mm, ChromXP C18-CL, 3 µm, 120 Å) with 100% A (0.1% formic acid in water, Fisher Chemicals, Geel, Belgium) for 10 min. The separation was performed at 300 nL/min, on a NanoLC column (Eksigent 75 µm x 15 cm, ChromXP 3C18-CL-120, 3 µm, 120 Å). The gradient was as follows: 0-1 min, 5% B (0.1% formic acid in acetonitrile, Fisher Chemicals, Geel, Belgium); 1-91 min, 5-30% B; 91-93 min, 30-80% B; 93-108 min, 80% B; 108-110 min, 80-5% B; 110-127 min, 5% B.

The samples used to generate the sequential window acquisition of all theoretical fragment ion spectra (SWATH)-MS spectral library were subjected to IDA using three different MS m/z ranges, which were calculated using the SWATH Variable Window Calculator V1.0 (Sciex, Framingham, US) based on a reference sample. The mass range for MS scan was set to m/z 400–614.9, 613.9–791.7, and 790.7–2,000. The MS/MS scan mass range was uniformly set to m/z 150–1,800. Following a TOF-MS survey scan (250 msec accumulation time), the 50 most intense precursors were selected for subsequent fragmentation and the MS/MS were acquired in high sensitivity mode for 40 msec, for a total cycle time of 2.3 s. The selection criteria for parent ions included an intensity of greater than 125 cps and a charge state ranging from +2 to +5. Once an ion had been fragmented through MS/MS, its mass was excluded from further MS/MS fragmentation for 12 s. The ions were fragmented in the collision cell using rolling collision energy, and CES was set to 5. Biological replicates (individual animals) were pooled to perform the IDA runs. Therefore, the 24 IDA MS raw files were combined and subjected to database searches in unison using ProteinPilot software v. 5.0 (Sciex, Framingham, US) with the Paragon algorithm to generate the Spectral Library. A UniProt reviewed database (17,032 entries, accessed on 16/10/2019) containing the sequences of Mus musculus was used. The following search parameters were set: Cys alkylation: Iodoacetamide; Digestion: Trypsin; Instrument: TripleTOF 6600; ID focus: Biological modifications and Amino acid substitutions; Search effort: Thorough; FDR analysis: Yes. Only the proteins with <1% FDR were considered.

##### Protein quantification by SWATH-MS

Five biological replicates from each condition were analysed by SWATH-MS, using the instrument setup described for the IDA runs. The mass spectrometer was set to operate in cyclic data-independent acquisition (DIA), similarly to the previously established method (Gillet et al., 2012). SWATH-MS data were acquired using the SWATH acquisition method, applying a set of 64 overlapping variable SWATH windows covering the precursor mass range of 400–2,000 m/z. The variable SWATH windows were calculated using the SWATH Variable Window Calculator V1.0 (Sciex, Framingham, US) based on the same reference sample. At the beginning of each cycle, a 10 ms survey scan (400-2,000 m/z) was acquired, and the subsequent SWATH windows were collected from 150 to 1,800 m/z for 50 ms, resulting in a cycle time of 3.26 s. The collision energy for each window was set using rolling collision energy, and CES was set to 5.

Data processing was performed using the SWATH processing plug-in for PeakView 2.2 (Sciex, Framingham, MA USA). In brief, peptides were selected from the library using the following criteria: i) the unique peptides for each protein were ranked by the precursor ion intensity from the IDA runs as estimated by the ProteinPilot software: ii) shared peptides (peptides that shared the same amino acid sequence between different protein entries) were excluded from selection. Up to 6 peptides were chosen per protein, and SWATH quantification was attempted for all proteins in the library file that were identified below 1% Global FDR from fit from ProteinPilot searches (which corresponded to a peptide confidence threshold of 98%). Target fragment ions, up to 6, were automatically selected. Manual inspection was performed for random peptides to check the quality of the auto selection and fragment ions were edited accordingly. Peak group confidence threshold was determined based on an FDR analysis using the target-decoy approach, and extraction FDR threshold was set to 1%. Peptides that met the 1% FDR threshold in all the five replicates were retained. The peak areas of the fragment ions were extracted using an XIC window of 6 minutes, and an XIC width of 20 ppm. Data were directly exported to Markerview 1.3.1 (Sciex, Framingham, MA USA) and normalized using total area sums to obtain the final quantification values. MarkerView was also used to perform the PCA and t-test statistical tests.

##### Quantitative analysis

An outlier removal approach was executed using GraphPad Prism Version 9, using iterative Grubb's using Alpha = 0.2. Missing values were replaced using average values from the corresponding group of samples. Protein intensities were then divided by the number of theoretically observable peptides between 6 and 30 amino acids, neglecting missed cleavages, to obtain normalized intensity-based absolute quantification (iBAQ) values[16]. The log2(x + 1) transformed iBAQ values were quantile normalization using the R function "normalize.quantiles"[17]. The normalized values were subjected to statistical analysis utilizing R package limma[18] where different contrast were specified. Correction for multiple testing was applied using the method of Benjamini & Hochberg[19].

#### Immunoblot analysis

##### Gel separation or dot-blots

10 or 15 µg of total protein from tissue lysates were separated by SDS-PAGE electrophoresis using a Tetra cell (Bio-Rad; Hercules, CA, USA), in 12% polyacrylamide separation gel and a 4% polyacrylamide stacking gel, applying a constant voltage of 120 V. Pre-stained standard proteins and pool sample, composed by 10 µl of cerebellum samples from each mouse, were also loaded onto the gel for inter-gel normalization purposes. 10 µg of total protein from tissue lysates were also loaded onto nitrocellulose membranes using a dot-blot system, and the wells washed twice with PBS before removing the membrane from the apparatus.

##### Western-blotting procedures

Gel separated proteins were transferred to nitrocellulose membranes, using standard procedures with a Mini Trans-Blot system (Bio-Rad; Hercules, CA, USA). Membranes were incubated with blocking solution (5% bovine serum albumin) in 1x TBS (20 mM Tris, 136 mM NaCl, pH 7.6) at room temperature for 30 minutes. Primary antibody incubations were carried out overnight at 4ºC, using given concentrations in blocking solution: $N^{\varepsilon}$-carboxyethyl lysine (CEL) (Mouse Anti- $N^{\varepsilon}$-carboxyethyl lysine, in a dilution of 1:1000 in blocking solution, Cosmo-Bio, USA), aSyn (Purified Mouse Anti-α-Synuclein antibody, in a dilution of 1:1000 in blocking solution, BD Biosciences; San Jose, CA, USA), and β-actin (Mouse Monoclonal anti-β-actin antibody in a dilution of 1:5000 in blocking solution, Ambion, Thermo Fisher Scientific; Waltham, MA, USA). Membranes were washed and incubated with secondary antibody (ECL™ Anti-Mouse IgG, HRP-Linked antibody in a dilution of 1:5000 in blocking solution, Amersham™; Little Chalfont, UK) for 1.5 hours. Detection procedures were carried on according to ECL system (GE Healthcare, Life Sciences; Little Chalfont, UK), and the signal detected using a ChemiDoc™ Imaging Systems (Bio-Rad, Hercules, CA, USA) with the most appropriate exposure time. Densitometry was performed using ImageJ - Image Processing and Analysis in Java[20]. When required, membranes were incubated with stripping solution (250 mM Glycine, 0.1 % of 10 % SDS, pH 2.0) for 45 minutes at room temperature with agitation, followed by 4 washing steps, twice with 1x TBS and twice with 1x TBS supplemented with 10% Tween 20 solution. Membranes were then incubated in blocking solution for 30 minutes before reprobing with the required antibodies.

### Supplementary Figures

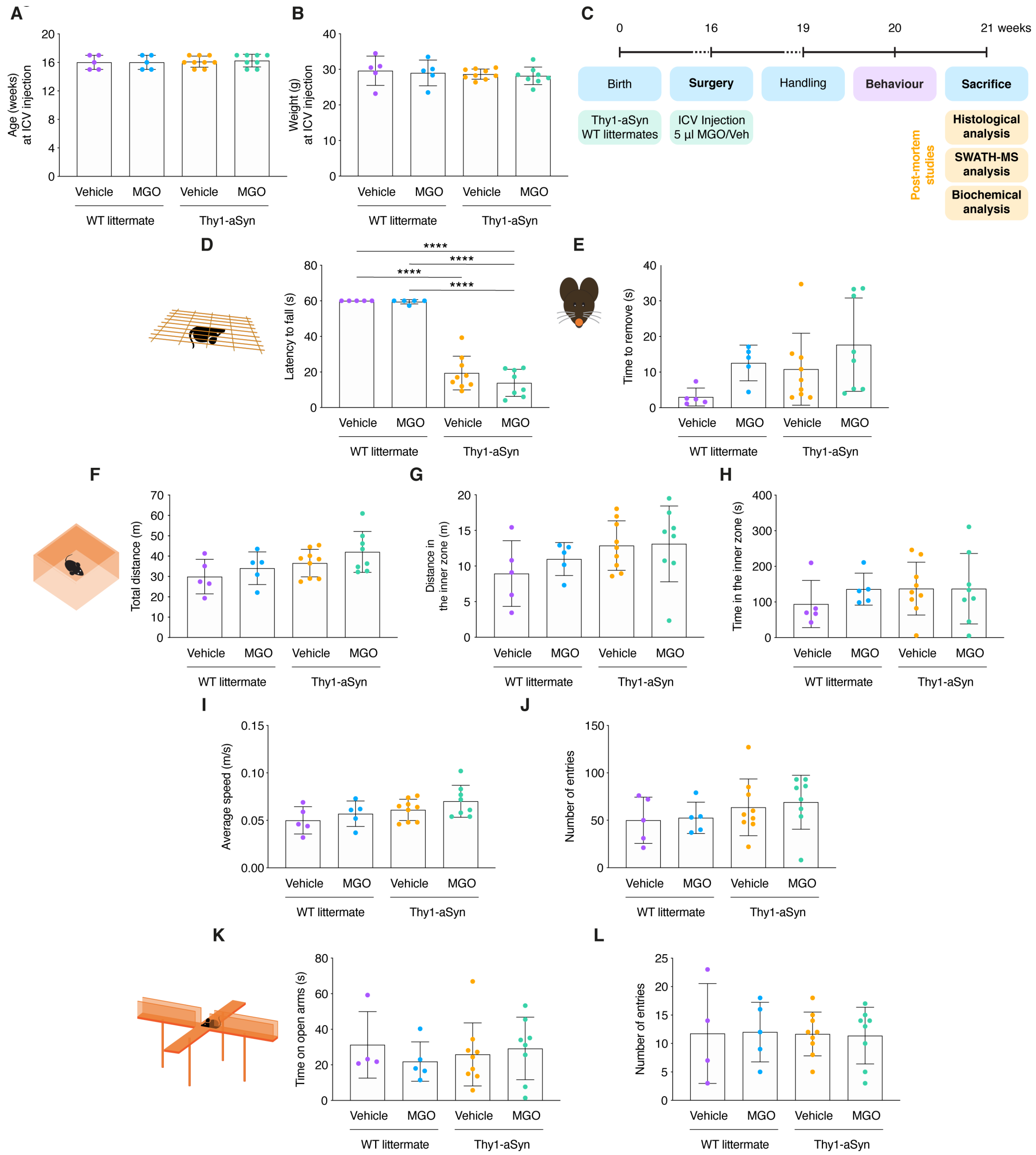


Supplementary Figure 1. Non-altered mice behaviour. **(A)** Schematic of experimental design: wild-type littermate (WT) and Thy1-aSyn transgenic (Tg) mice received an intracerebroventricular (ICV) injection of MGO or vehicle (PBS) at 16 weeks of age. Behavioural testing started 4 weeks post-surgery. Plot representations of mice demographics: **(B)** age in weeks and **(C)** weight in g at MGO ICV injection. Behavioural testing was performed 4 weeks after surgery. Plot representations of: **(D)** wire hang test; **(E)** adhesive removal test; open field test - **(F)** total distance, **(G)** distance in the inner zone, **(H)** time in the inner zone, **(I)** average speed, **(J)** number of entries; Elevated plus maze test - **(K)** time on open arms, and **(L)** number of entries; At least n=5 in all groups.

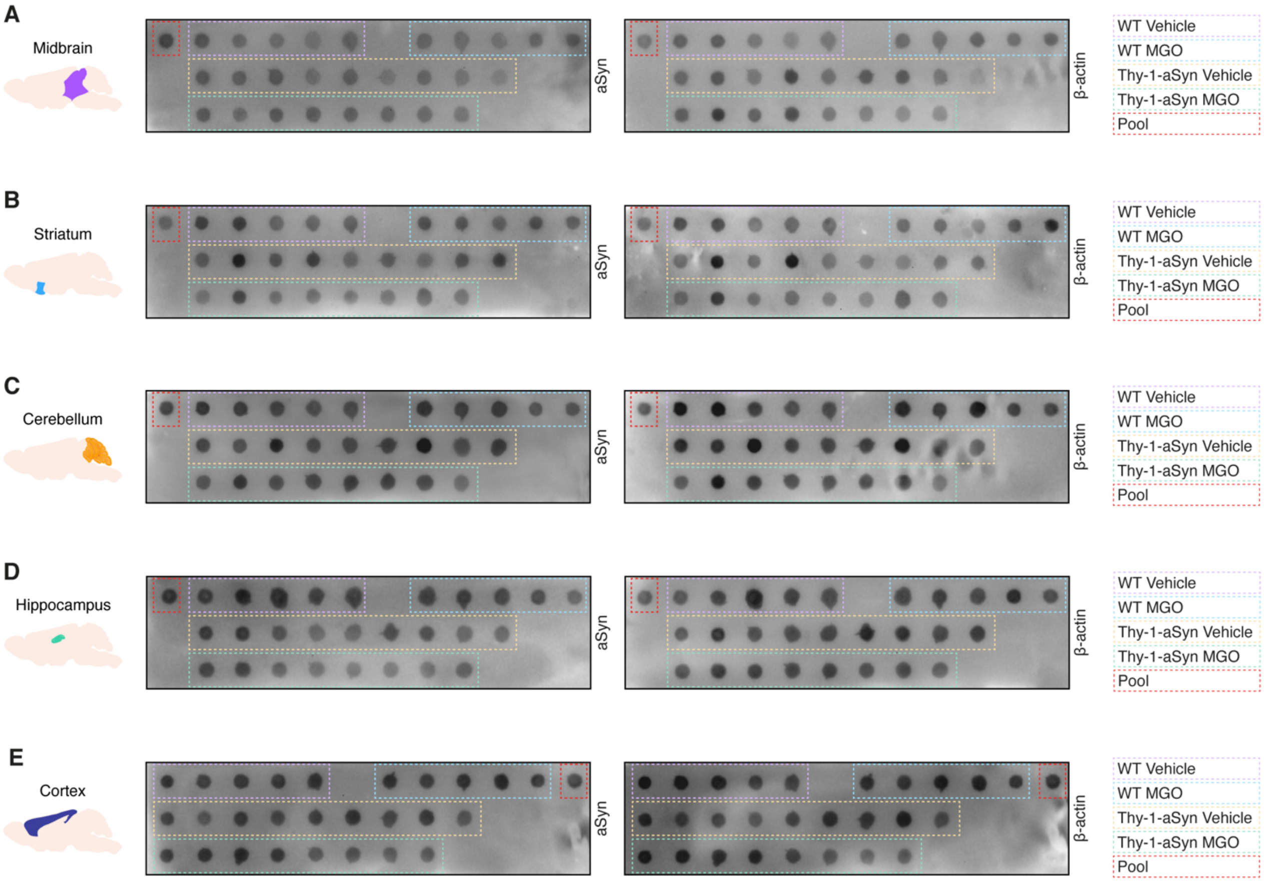


**Supplementary Figure 2. Signal of glycated protein in different brain regions of all experimental groups.** Wild-type littermate (WT) and Thy1-aSyn transgenic mice received an intracerebroventricular (ICV) injection of MGO or vehicle (PBS) and protein brain extracts from several regions analysed 5 weeks post-injection. Protein extracts loaded into nitrocellulose membranes in a dotblot system and probed with anti-CEL and anti-β-actin, for normalization. Representative blots from all samples from each experimental group and a pool sample for inter-gel comparison are shown for **(A)** midbrain, **(B)** striatum, **(C)** cerebellum, **(D)** hippocampus, and **(E)** frontal cortex, respectively.

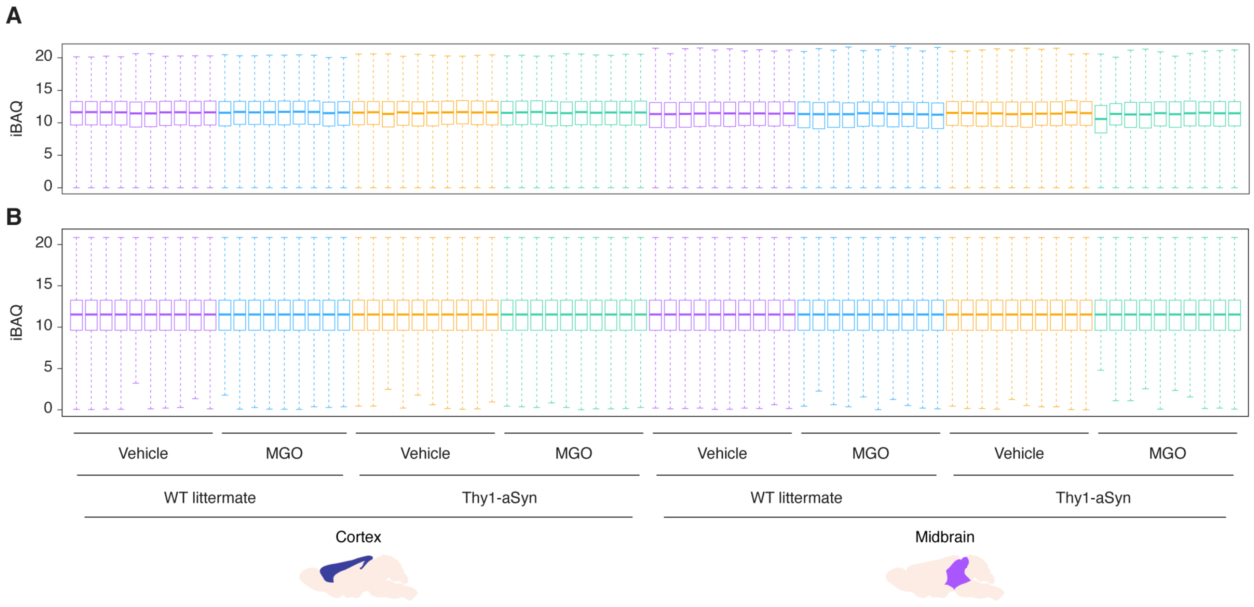


Supplementary Figure 3. Boxplots of intensity-based absolute quantification (iBAQ) values of the SWATH-MS proteomic analysis. Boxplot representation of outlier-removed **(A)** raw and **(B)** normalised iBAQ values for each performed SWATH-MS run.

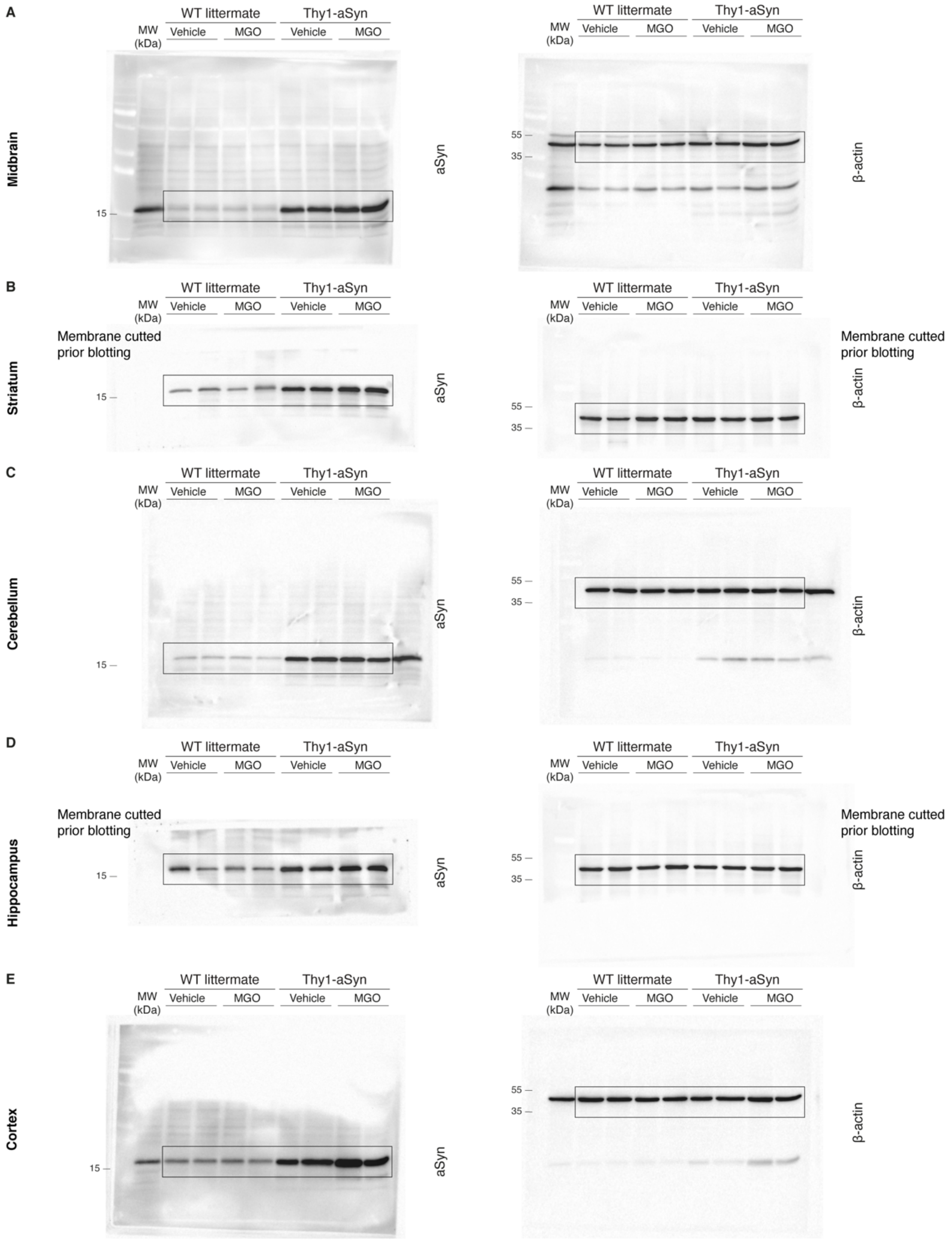


Supplementary Figure 4. Uncropped blots of main Figure 2. Representative blots showing 2 samples from each experimental group are shown for aSyn probing: **(A)** midbrain, **(B)** striatum, **(C)** cerebellum, **(D)** hippocampus, and **(E)** prefrontal cortex.
